# Neurogranin modulates the Rate of Association between Calmodulin and Target Peptides

**DOI:** 10.1101/2024.03.21.586151

**Authors:** John A. Putkey, Laurel Hoffman, Vladimir Berka, Xu Wang

## Abstract

The best-known mode of action of calmodulin (CaM) is binding of Ca^2+^ to its N- and C-domains, followed by binding to target proteins. An underappreciated facet of this process is that CaM is typically bound to proteins at basal levels of free Ca^2+^, including the small, intrinsically disordered, neuronal IQ-motif proteins called PEP-19 and neurogranin (Ng). PEP-19 and Ng would not be effective competitive inhibitors of high-affinity Ca^2+^-dependent CaM targets at equilibrium since they bind to CaM with relatively low affinity, but they could influence the time course of CaM signaling by affecting the rate of association of CaM with high-affinity Ca^2+^-dependent targets. This mode of regulation may domain specific since PEP-19 binds to the C-domain of CaM, while Ng binds to both N- and C-domains. In this report, we used a model CaM binding peptide (CKIIp) to characterize the preferred pathway of complex formation with Ca^2+^-CaM at low levels of free Ca^2+^ (0.25 to 1.5 µM), and how PEP-19 and Ng affect this process. We show that the dominant encounter complex involves association of CKIIp with the N-domain of CaM, even though the C-domain has a greater affinity for Ca^2+^. We also show that Ng greatly decreases the rate of association of Ca^2+^-CaM with CKIIp due to the relatively slow dissociation of Ng from CaM, and to interactions between the Gly-rich C-terminal region of Ng with the N-domain of CaM, which inhibits formation of the preferred encounter complex with CKIIp. These results provide the general mechanistic paradigms that binding CaM to targets can be driven by its N-domain, and that low-affinity regulators of CaM signaling have the potential to influence the rate of activation of high-affinity CaM targets and potentially affect the distribution of limited CaM among multiple targets during Ca^2+^ oscillations.

**STATEMENT OF SIGNIFICANCE:** Calmodulin is a small, essential regulator of multiple cellular processes including growth and differentiation. Its best-known mode of action is to first bind calcium and then bind and regulate the activity of target proteins. Each domain of CaM has distinct calcium binding properties and can interact with targets in distinct ways. We show here that the N-domain of calmodulin can drive its association with targets, and that a small, intrinsically disordered regulator of calmodulin signaling called neurogranin can greatly decrease the rate of association of CaM with high-affinity Ca^2+^-dependent targets. These results demonstrate the potential of neurogranin, and potentially other proteins, to modulate the time course of activation of targets by a limited intracellular supply of calmodulin.

## INTRODUCTION

Calmodulin (CaM) is a highly conserved and ubiquitous Ca^2+^ sensor protein that is essential for the survival and function of all eukaryotic cells. The global structure of CaM consists of N- and C-terminal Ca^2+^ binding domains that are connected by a flexible linker. This simple design allows multiple modes of interaction with targets that can involve only one or both domains, and in the presence or absence of Ca^2+^ (1–4). The most familiar mode of interaction is high-affinity, Ca^2+^-dependent binding of both N- and C-domains of CaM to a relatively short and contiguous target sequence to form a compact structure that resembles two hands grasping a rope as illustrated in Figure 1.

**Figure 1:**
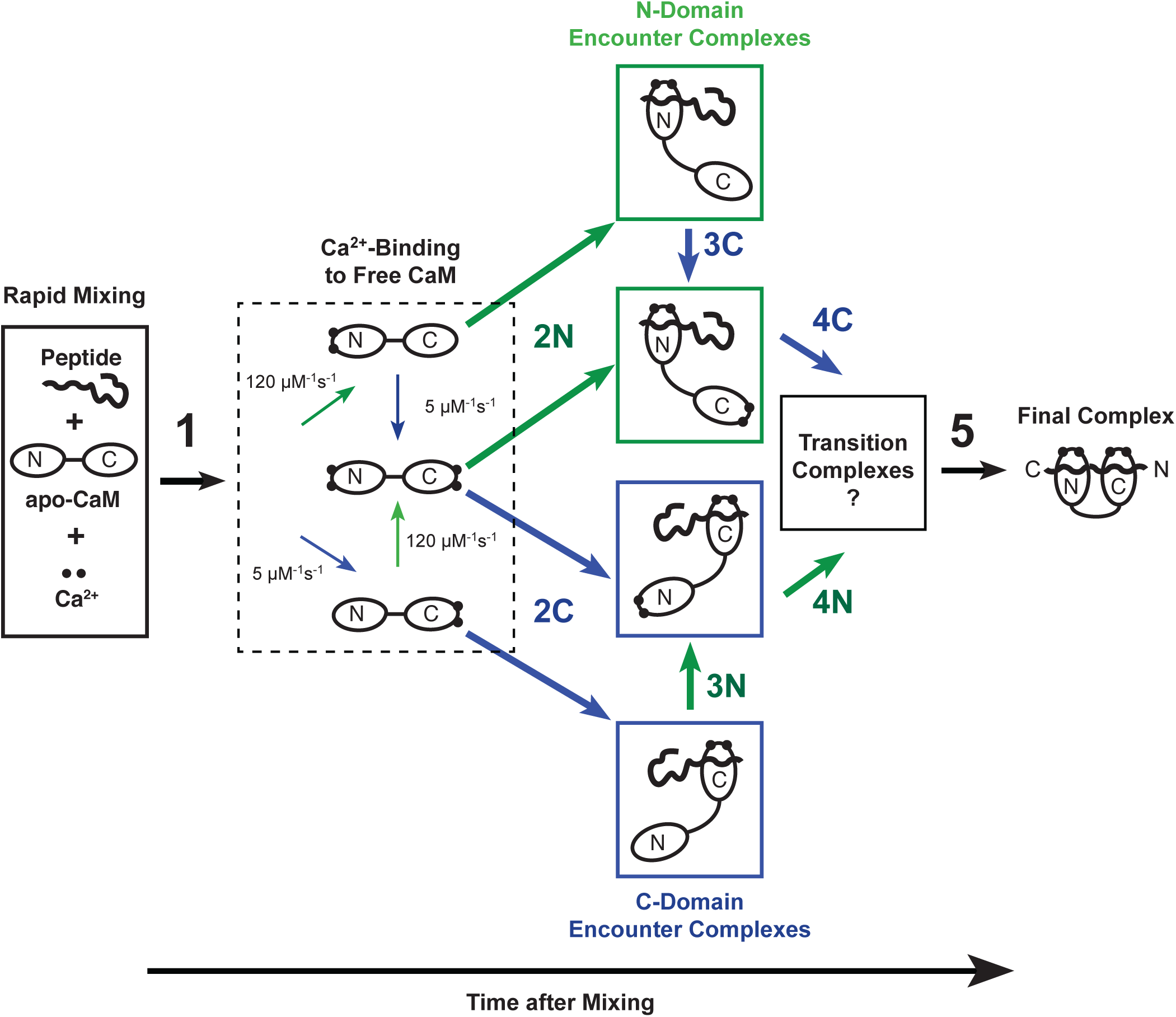
Pathways of interactions between Ca^2+^, CaM and a CaM binding peptide. Important parameters are described in the text. Green and Blue arrows indicate binding of Ca^2+^ or CKIIp to the N-domain or C-domain of CaM, respectively. The final complex has the peptide bound to CaM in an antiparallel orientation in which the N- and C-domains of CaM associate primarily with the C- or N-term of the CKIIp, respectively. However, a minor population of transient transition complexes may exist that have a parallel binding orientation.

Interactions between CaM and Ca^2+^-dependent target proteins are primarily regulated by the amplitude and frequency of oscillations in intracellular free Ca^2+^ levels (Ca^2+^). However, the distribution of CaM among its multiple targets during Ca^2+^ oscillations will depend on target-specific rates of association and dissociation with Ca^2+^-CaM, and how targets affect the Ca^2+^ binding properties of CaM. In general, there is an inverse relationship between the rate of dissociation of Ca^2+^ from CaM and the affinity of a target for CaM, which results in slower release of Ca^2+^ and higher affinity when CaM is bound to high affinity targets (5, 6). Less is known about the rate of association of Ca^2+^-CaM with target proteins, and if the N- or C-domain plays a dominate role. Since the N- and C-domains can interact separately with CaM binding sites on proteins (7), formation of a final active complex could proceed via multiple pathways that are initiated by Ca^2+^ binding to either the N- or C- domains as shown in Figure 1. Early models for association of CaM with binding peptides focused on formation of an initial encounter complex involving the C-domain of CaM along pathway 2C in Figure 1 (6, 7), primarily because the affinity of Ca^2+^ binding to the C-domain is greater than the N-domain (8). However, two factors suggest that the N-domain could play a dominant role in formation of the encounter complex along pathway 2N. First, the rate of Ca^2+^ association with the N-domain is a least 25-fold greater than the C-domain, and second the affinity of Ca^2+^ binding to both the N- and C-domains of CaM is greatly increased upon association with targets (5, 6, 9, 10).

Two other factors can impact the mechanism and rate of association of Ca^2+^-CaM with targets. The first is that levels of Ca^2+^_F_ can vary significantly in different subcellular compartments. Resting free Ca^2+^ is around 0.1 µM, and can increase 10-fold or greater in response to a variety of stimuli. Peak levels of free Ca^2+^ in the bulk cytoplasm may increase to 1 to 2 µM, while it may be up to 100 µM near the surface of membranes where Ca^2+^ is released through channels. Interactions between CaM and targets at lower levels of free Ca^2+^ is of special interest since the distinct Ca^2+^ binding properties of the N- and C-domains may have a greater potential to influence the mechanism of association. The other factor that could modulate interactions between Ca^2+^-CaM and its target are proteins that bind to apo-CaM. Two small intrinsically disordered IQ-motif proteins called PEP-19 and neurogranin (Ng) bind to either apo or Ca^2+^- CaM (11, 12). PEP-19 binds selectively to the C-domain of CaM (13), while Ng interacts both N- and C-domains (14). PEP-19 greatly increases the rates of Ca^2+^ association and dissociation at the C-domain without significantly affecting the K_Ca_ (13), which makes the kinetics of Ca^2+^ binding to the C-domain more similar to those of the N-domain. Ng also increases the rates of Ca^2+^ dissociation at the C-domain of CaM, but has a lesser effect on the Ca^2+^ association rate, thereby decreasing overall Ca^2+^ binding affinity (15). These properties of PEP-19 and Ng on CaM have the potential to regulate activation of Ca^2+^-dependent CaM target proteins by simple competition, or by modulating the rates of binding Ca^2+^ to CaM.

The goal of the current study was to use a novel experimental model system to first determine the kinetics and domain specificity of CaM binding to a model CaM binding peptide in response to levels of free Ca^2+^ between 0.2 µM and 5 µM, and then determine if PEP-19 and/or Ng affect the rates of peptide-binding. Our data show that the dominant pathway for association involves formation of an encounter complex between the N-domain of CaM and the binding peptide. PEP-19 has minimal effect on the rate of binding CaM to the model peptide, however, Ng greatly decreases the rate of association and this effect is most pronounced at the lower levels of free Ca^2+^. The effects of Ng are due to its ability to bind to the N-domain of CaM, which likely inhibits formation of the dominant encounter complex. These data demonstrate that Ng could have broad effects on CaM signaling pathways by modulating the rate of association of CaM with Ca^2+^-dependent targets.

## MATERIALS AND METHODS

### Protein and peptides

Wild type mammalian calmodulin (CaM), CaM(K75C), neurograinin (Ng), and Purkinje cell protein 4 (PEP-19) were expressed and purified as previously described (13, 15, 16). Peptides Ng(26–49), comprising the IQ domain (ANAAAAKIQASFRGHMARKKIKSG), and Ng(13–49), comprising the IQ domain and adjacent acidic region (DDDILDIPLDDPGANAAAAKIQASFRGHMARKKIKSG), were synthesized by LifeTein. The peptide CKIIp, representing the CaM binding site of CaM kinase II (FNARRKLKGAILTTMLATTRN), and a modified version containing a C-terminal cysteine for fluorescent labeling (FNARRKLKGAILTTMLATTRNGC), were synthesized by GenicBio Limited. Purchased peptides were resuspended in 0.13% TFA and further purified on a reverse phase C18 column (Waters) using a gradient of acetronitrile, 0.1% TFA. Fractions containing peptides were lyophilized and resuspended in water.

Buffers used for stopped flow experiments were decalcified by passage over a small column of Calcium Sponge (Molecular Probes/ThermoFisher). Indo-1 was used to determine the effectiveness of decalcification. CaM was decalcified using EDTA/BAPTA followed by desalting. Typically, 20 to 50 mg of CaM in 10 ml of buffer was made 2 to 5 mM in EDTA and 0.1 mM BAPTA, which was used a chromophore to monitor desalting. The solution was desalted using a P6DG column in a mobile phase of 20 mM ammonium bicarbonate, pH 8.0 that was decalcified using Calcium Sponge. Effective separation of CaM from the chelators was monitored by UV absorbance. The decalcified CaM was lyophilized and weighed.

CaM(K75C) was labeled with the fluorescent probe 5-((((2-iodoacetyl)amino)ethyl)amino)naphthalene-1-sulfonic acid (1,5-IAEDANS from Molecular Probes) as previously described (17) to yield CaMD. Donor/acceptor-labeled CaMD/A was prepared by labeling T34C/T110C CaM with the donor 1,5-IAEDANS and the chromophoric acceptor N-[4-(dimethylamino)-3,5-dinitrophenyl]-maleimide (DDPM) as described previously (18). CKIIp with a C-terminal cysteine was labeled with DDPM to yield CKIIpA. CKIIp in 40 mM MOPS, pH 8.0, was first reduced with with 2.5 mM TCEP for 30 minutes, then reacted with 1.1 molar excess DDPM for 4 hours at room temperature. The labeled peptide was purified on a C18 column, lyophilized, and resuspended in water. Labeling of Ng with DDPM (NgA) was carried out in a similar fashion.

### Assay for the rate of association between CaM and a CKIIp

Several experimental systems were employed to determine the rate of association of CaM and CKIIp in the absence or presence of PEP-19 or Ng. The first system used changes in Tyr fluorescence that results from Ca^2+^-dependent conformational changes in the C-domain of CaM. Unlabeled CaM was excited at 280 nm and emission was monitored from 300-400 nm. The two other systems used FRET. The inter-FRET assay had the donor on CaMD and acceptor on CKIIpA, while the intra-FRET assay had the donor and acceptor on CaMD/A. IAEDANS-labeled proteins were excited at 340 nM, and FRET was monitored as a decrease in fluorescence 495 nm.

All rates were measured in an Applied Photophysics Ltd model SV.17 MV sequential stopped-flow spectrofluorimeter with an instrument dead time of 1.7 ms. Reactions were initiated by rapidly mixing equal volumes of solutions from syringes A and B. The stopped-flow buffer (SFB) contained 20 mM MOPS, pH 7.5, 100 mM KCl and 1 mM DTT. Syringe A contained SFB with 2.5 µM CaM with 10 µM BAPTA added to ensure that CaM was in the apo state. Concentrations of PEP-19 or Ng in syringe A were 40 µM and Ng 20 µM, respectively, to maximize binding to apo CaM given that Ng binds to CaM with higher affinity that PEP-19, and to give the same ratios of CaM:CKIIp:PEP-19 or Ng as in the NMR experiments shown in Supplemental Figures 2 to 4. Syringe B contained SFB with 5 µM CKIIp, 5 mM HEDTA and sufficient CaCl_2_ to achieve a desired free Ca^2+^ (Ca^2+^_F_) based on Maxchelator Standard (available at http://maxchelator.stanford.edu/webmaxc/webmaxcS.htm). The actual Ca^2+^_F_ was then verified experimentally using 5,5’-Dibromo BAPTA (Br_2_BAPTA) absorbance at 263 nm in a UV-VIS spectrometer using the following equation:

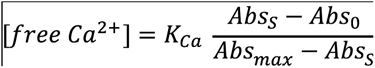

where the K_Ca_ for Br_2_BAPTA is 1.59 μM, Abs_S_ is the absorbance of the sample, Abs_0_ is absorbance in the absence of Ca^2+^ and Abs_max_ is the absorbance in the presence of saturating 5 mM CaCl_2_.

A key requirement of the assay is that binding of Ca^2+^ to CaM should not affect the equilibrium levels of Ca^2+^ in the buffered solutions. To test this, 5 µM Br_2_BAPTA was added to the Ca^2+^/HEDTA as a marker for free Ca^2+^ levels and this was then mixed with 10 µM EDTA to mimic 4 Ca^2+^ binding sites in 2.5 µM CaM. The absorbance of Br_2_BAPTA was unaffected, which indicated that the Ca^2+^ was not significantly changed by binding Ca^2+^ to EDTA. A second requirement is that the rate of replenishing the pool of Ca^2+^ as Ca^2+^ binds to CaM should not be a rate-limiting step. To confirm this requirement, we used the Ca^2+^ chromophore BAPTA, which binds Ca^2+^ with a K_d_ of 0.16 µM and k_on_ of 450 µM^-1^s^-1^. If replenishing the pool of Ca^2+^ is not rate limiting, then 10 µM BAPTA should be saturated with Ca^2+^ within 10 ms after mixing with 5 mM HEDTA buffers with Ca^2+^ set at 1-5 µM. Control experiments using BAPTA showed that replenishing the pool of Ca^2+^ is not rate-limiting in our experimental system.

Time-dependent changes in fluorescence were fit to the following equations with single, double or triple exponential terms:

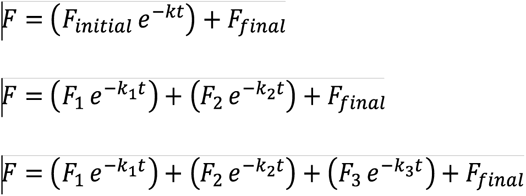

Where F is the observed fluorescence intensity at time t, F_initial_ is fluorescence at t=0, F_final_ is the final fluorescence and k is the observed rate.

### CaM-Ng dissociation rate

The rate of dissociation of Ng from Ca^2+^-CaM was determined with stopped flow experiments using CaM(D2C) labeled with IAEDANS and Ng labeled with DDPM (NgA) at its endogenous Cys residues. Syringe A contained 20 mM MOPS, pH7.5, 100 mM KCl, 5 mM CaCl_2_ 1 μM IAEDANS-labeled CaM(D2C), 10 μM NgA, while syringe B contained 20 mM MOPS, pH7.5, 100 mM KCl, 5 mM CaCl_2_ and 20 μM unlabeled CaM. The decrease in FRET fluorescence was monitored as unlabeled CaM exchanged with labeled CaM.

### NMR methodology

All NMR experiments were performed on a DRX 600 MHz spectrometer equipped with a 5 mm triple-resonance cryoprobe at 310 K. NMR buffer contained 10 mM imidazole, 100 mM KCl, 5 mM CaCl_2_, 0.1 mM [^15^N]-CaM, 5 mM DTT and 5% D_2_O at pH 6.3. When present, CKIIp, Ng and PEP-19 were added at concentrations of 0.2, 0.8 and 1.6 mM respectively. NMR spectra were collected using 4 scans and a spectral width of 32 ppm in the 15N dimension. Backbone assignments for free Ca^2+^-CaM were reported previously (13).

All NMR data were processed and analyzed using Bruker Topspin 3.5. ^1^H chemical shifts were referenced to DSS (2,2-dimethyl-2-silapentane-5-sulphonate). The average amide chemical shift change was calculated using the following equation:

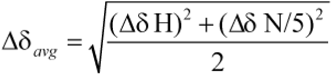

where Δδ_H_ is the change of ^1^H chemical shift and Δδ_N_ is the change of ^15^N chemical shift.

## RESULTS

### Dissociation of Ca^2+^ from CaM and the CaM/CKIIp complex

We used a Ca^2+^-dependent CaM binding peptide (CKIIp) derived from a.a. 293-312 of CaM-dependent kinase II as a model system. The structural features of binding CKIIp to Ca^2+^-CaM are known in detail from x-ray crystallography (19). The peptide can be thought of as a bivalent CaM binding site since a single peptide binds to both the N- and C-domains of Ca^2+^-CaM. It has a 1-5-10 spacing of hydrophobic residues with Leu-299 and Leu-308 anchoring the peptide to the C- and N-domains of CaM, respectively, via hydrophobic interactions (3, 19). CKIIp binds to Ca^2+^-CaM with very high affinity (Kd = 1.6 pM), and with a fast k_on_ of 1.2 x 10^8^ M^-1^s^-1^ (16). This makes CKIIp a stringent test for the ability of PEP-19 and Ng to modulate the rate of association of a target with Ca^2+^-CaM.

To validate our various assays, and to provide a complete set of kinetic parameters, we first determined the rates of dissociation of Ca^2+^ and CKIIp from CaM. Fluorescence from Tyr-99 and Tyr-138 is a well-established marker for Ca^2+^-dependent events in the C-domain of CaM. Figure 2A and Table 1 show the rate of change in Try florescence after free Ca^2+^-CaM is rapidly mixed with excess EGTA. A single rate of 9.2 sec-1 was observed, which corresponds well to the rate of dissociation of Ca^2+^ from sites III and IV of CaM reported in previous studies (6, 13). The Ca^2+^ sensitive fluorescent dye Quin-2 was also used since it is a direct measure of Ca^2+^ release and can be expressed as moles Ca^2+^ released per mole of CaM. Figure 2B and Table 1 show that a biphasic release of Ca^2+^ from free CaM was observed using Quin-2. The majority of the fast phase of Ca^2+^ release occurs in the dead time of the stop flow (2 ms). The rate was estimated to be about 1300 s-1 (see Table 1) and is due to release of Ca^2+^ from the N-domain. The slow phase of 8.9 s-1 corresponds well to the rate measured using Tyr fluorescence and is consistent with earlier results for Ca^2+^-dissociation from the C-domain. Nearly identical rates for Ca^2+^ release measured using both Try and Quin-2 demonstrates that the rate of conformational relaxation in the C-domain is tightly coupled with Ca^2+^ dissociation.

**Figure 2:**
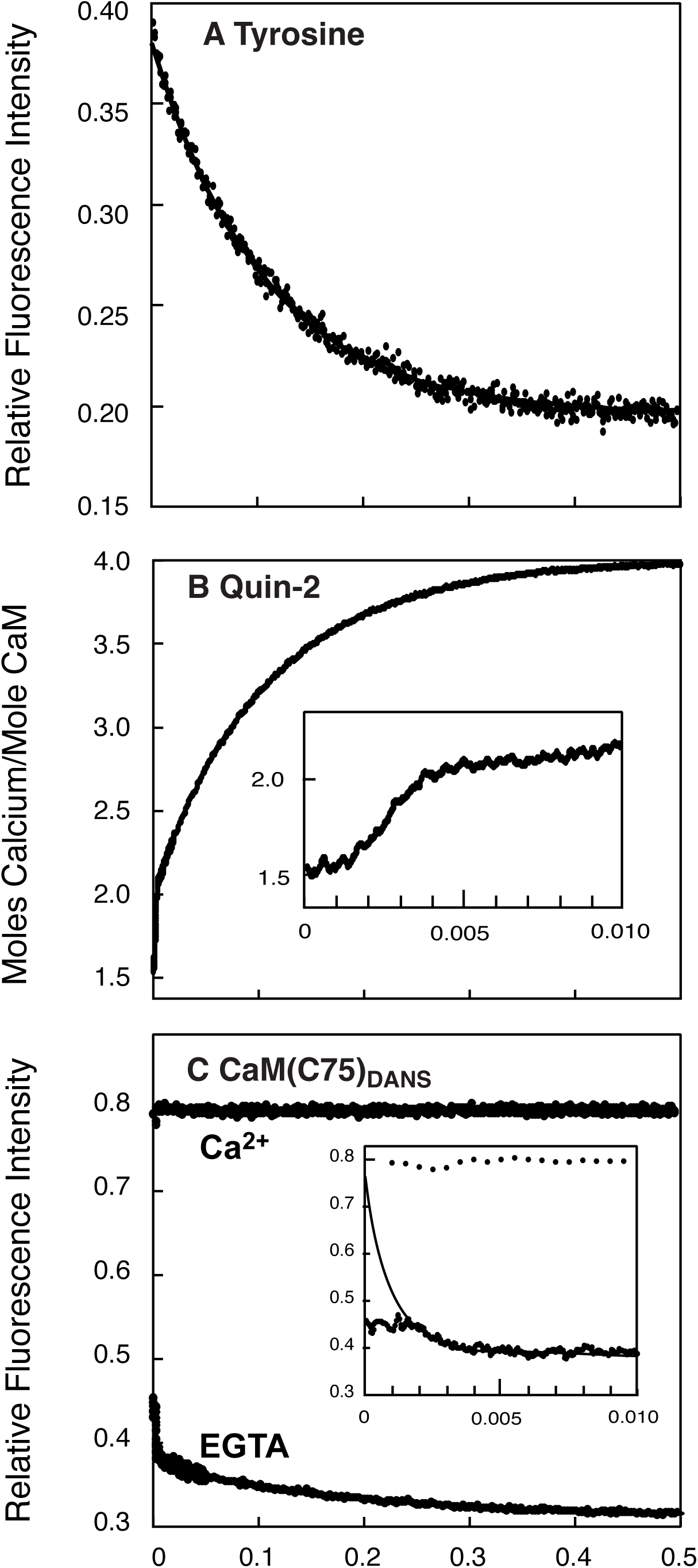
Rate of dissociation of Ca^2+^ from free Ca^2+^-CaM. Ca^2+^ dissociation was detected by intrinsic Tyr fluorescence (*Panel A*), the fluorescent Ca^2+^ chelator Quin-2 (*Panel B*), or CaMD (*Panel C*). Syringe A contained 5 µM CaM or CaMD and 200 µM CaCl_2_. Syringe B contained 5 mM EGTA (*Panels A* and *C*) or 400 µM Quin-2 (*Panel B*). The Quin-2 response was calibrated using an Orion Ca^2+^ standard. The slow phase of Ca^2+^ release in *Panel B* corresponds to 2 moles Ca^2+^ released/mole CaM, and could be used as a calibration control. *Panel C* shows a control experiment (upper data set) in which buffer B containing 200 μM CaCl_2_ rather than 5 mM EGTA. This provides an initial fluorescence value used to fit the fast phase of fluorescence shown in the inset of *Panel C* .

**Table 1.** Rates of Dissociation of ca^2^+ from Ca^2^+-caM in the Absence or Presence of CKIIp.

|  | Free Ca<sup>2+</sup>-CaM |  |  |  | Ca<sup>2+</sup>-CaM + CKIIp |  |  |  |
| --- | --- | --- | --- | --- | --- | --- | --- | --- |
|  | Rate 1 | Amp 1 | Rate 2 | Amp 2 | Rate 1 | Amp 1 | Rate 2 | Amp 2 |
| Quin-2 | 1300 ± 120 | 2 | 8.9 ± 0.1 | 2 | 6.3 ± 1 | 2.1 | 0.27 ± 0.04 | 2 |
| Tyrosine | --- | --- | 9.2 ± 0.2 | 1 | 5.5 ± 0.1 | 0.17 | 0.25 ± 0.01 | 0.83 |
| CaM(C75D) | 1600 ± 140 | 0.86 | 5.2 ± 0.2 | 0.14 | 5.9 ± 0.1 | 0.79 | 0.34 ± 0.01 | 0.21 |
| FRET |  |  |  |  | 5.8 ± 0.8 | 0.37 | 0.23 ± 0.02 | 0.63 |

All rates are the average ± SD of 4-9 independent experiments.
The amplitudes for data collected using Quin-2 indicate number of moles Ca<sup>2+</sup> released/mole protein and were obtained by calibrating the response of Quin-2 with Ca<sup>2+</sup> standards. The moles Ca<sup>2+</sup> released/mole protein for the fast phase of free CaM could not be accurately fit since about 75% of the signal occurs during the dead time of the instrument. The value of 2 was assigned since CaM binds 4 moles Ca<sup>2+</sup>, and 2 moles Ca<sup>2+</sup>/mole protein were released during the slow phase.
The average amplitudes are given. Amplitudes for Tyr, CaM(C75D) and FRET indicate the relative fractional amplitude of each phase with 1.0 defined as the total change in fluorescence. Absolute values are used to determine amplitudes for FRET.
---- indicates that the best fit was a single exponential

**Table 2.** Rates of Association of CKIIp with Ca^2+^-CaM in the Absence of PEP-19 or Ng.

| Ca<sup>2+</sup><br>(μM) | Tyrosine Fluorescence | | | | | | Inter-FRET | | | | Intra-FRET | |
| --- | --- | --- | --- | --- | --- | --- | --- | --- | --- | --- | --- | --- |
|  | Free CaM |  | CaM +CKIIp |  |  |  |  |  |  |  |  |  |
|  | Rate<br>(s<sup>-1</sup>) | Amp | Rate 1<br>(s<sup>-1</sup>) | Amp 1 | Rate 2<br>(s<sup>-1</sup>) | Amp 2 | Rate 1<br>(s<sup>-1</sup>) | Amp 1 | Rate 2<br>(s<sup>-1</sup>) | Amp 2 | Rate<br>(s<sup>-1</sup>) | Amp |
| 0.26 | nd | | 1.5 ± 0.1 | 50 | 0.45 ± 0.01 | 49 | 13 ± 0.2 | 66 | 2.4 ± 0.1 | 26 | 1.2 ± 0.2 | 87 |
| 0.50 | 8.2 ± 0.5 | 9 | 5.0 ± 0.2 | 45 | 1.51 ± 0.03 | 56 | 26 ± 0.3 | 76 | 4.0 ± 0.1 | 20 | 3.2 ± 0.3 | 95 |
| 1.10 | 11.4 ± 0.3 | 27 | 12.9 ± 0.4 | 50 | 4.75 ± 0.21 | 50 | 68 ± 0.5 | 80 | 8.2 ± 0.2 | 20 | 10.1 ± 0.5 | 100 |
| 2.19 | 17.8 ± 0.3 | 56 | 22.7 ± 0.8 | 93 | | | 163 ± 2.6 | 86 | 15.9 ± 0.4 | 14 | 30.2 ± 2.3 | 95 |
| 3.26 | 30.7 ± 0.5 | 68 | 43.2 ± 0.8 | 93 | | | 278 ± 2.5 | 85 | 27.2 ± 1.0 | 14 | 49.2 ± 2.5 | 86 |
| 4.56 | 46.5 ± 0.7 | 84 | 69.9 ± 0.7 | 87 | | | 377 ± 2.2 | 74 | 36.1 ± 1.2 | 16 | 73.1 ± 2.2 | 74 |

At least 5 stopped-flow traces were averaged and fitted using AppliedPhotophysics SX-Pro Data. The rates are shown with the estimated error of the fit.
To allow comparisons of amplitudes obtained from biphasic fits, as well as fits for different levels of Ca<sup>2+</sup><sub>F</sub>, the amplitudes are expressed relative to the overall amplitude at the Ca<sup>2+</sup><sub>F</sub> that gave the largest overall change in fluorescence within the data set. In the absence of CKIIp, the largest change in fluorescence was at 5 mM Ca<sup>2+</sup>.

We also used IAEDANS-labeled CaM(K75C) (CaMD) to measure rates of Ca^2+^ release from free CaM. We showed previously that steady state fluorescence from CaMD was sensitive to conformational changes caused by Ca^2+^ binding and subsequent binding to CKIIp (20). Figure 2C and Table 1 show a decrease in fluorescence with a rate of approximately 1,600 s-1 occurs after rapidly mixing Ca^2+^-CaMD with excess EGTA, followed by a second phase with a smaller amplitude and slower rate of 5.2 s-1. We interpret the fast and slow phases to correspond to release of Ca^2+^ from the N- and C-domains, respectively. The larger amplitude for the fast phase is consistent with the location of C75 in helix D near the N-domain of CaM.

We next used Quin-2, CaMD, and Tyr fluorescence to determine how CKIIp binding to Ca^2+^-CaM affects the rates of Ca^2+^dissociation. Figure 3 and Table 1 show that dissociation of Ca^2+^ from the CaM/CKIIp complex was biphasic and that all rates were greatly reduced relative to free CaM. The fast rate was 5.5 to 6.3 s-1, while the slow rate was 0.27 to 0.34 s-1. These values correspond well with rates reported previously for dissociation of Ca^2+^ from CaM bound to CaM kinase II peptides (5, 6). We can confidently assign the fast and slow rates to dissociation of Ca^2+^ from the N- and C-domains, respectively, based on the amplitudes of responses. First, it is reasonable to conclude that the slow rate with the large amplitude (0.83) measured by Try fluorescence is due to conformational changes in the C-domain, while the faster rate with a much smaller amplitude (0.17) is due to an allosteric effect relayed through CKIIp to the C-domain when Ca^2+^ dissociates from the N-domain. Second, the fast rate observed using CaMD is associated with the largest amplitude (0.79), and this amplitude is comparable in magnitude to that observed due to dissociation of Ca^2+^ from the N-domain of free CaM. Similarly, the amplitudes of the slow rates observed using CaMD are similar in magnitude for both free CaM and the CaM/CKIIp complex.

**Figure 3:**
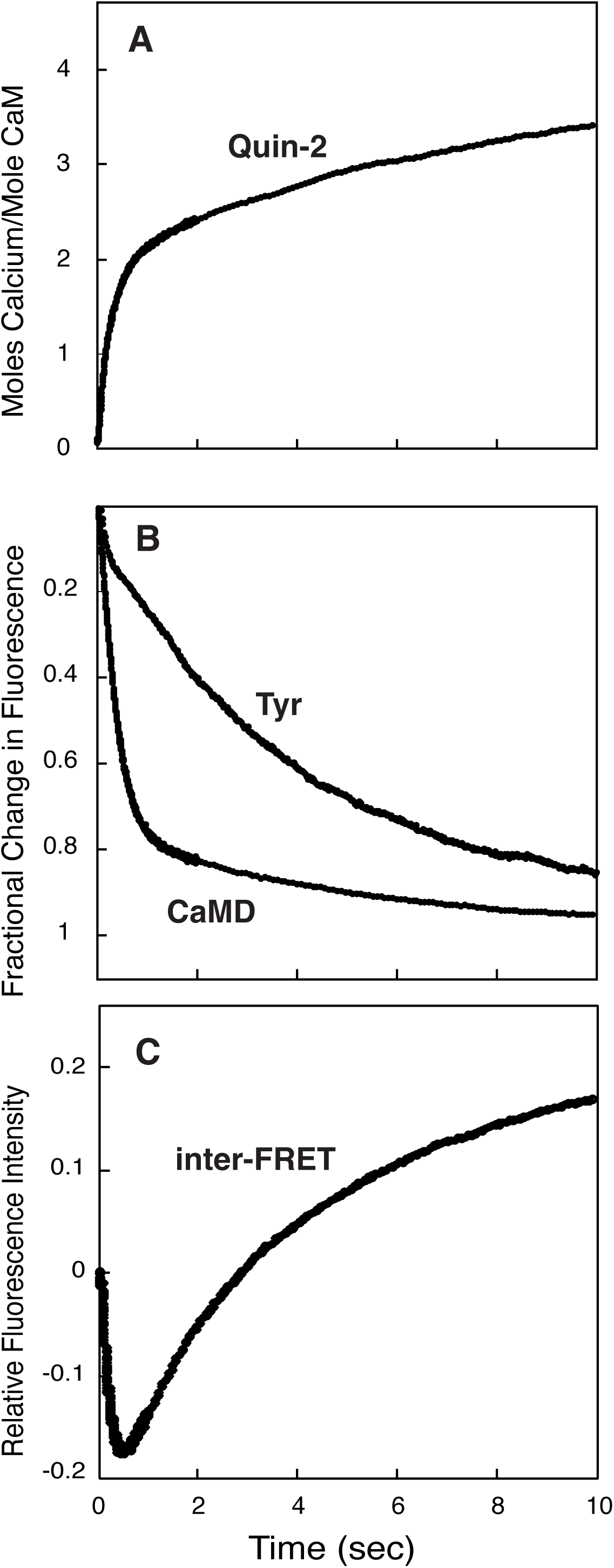
Rate of dissociation of Ca^2+^ from the Ca^2+^/CKIIp complex. *Panel A* shows the release of Ca^2+^ from the Ca^2+^/CKIIp complex detected using Quin-2. *Panel B* monitors conformational changes due to release of Ca^2+^ using fluorescence from Tyr or IAEDANS-labeled CaMD. *Panel C* uses a FRET assay to detect changes in distance between CaM and CKIIp after Ca^2+^ is removed. The FRET donor is IAEDANS-labeled CaM(C110)D, and the acceptor is DDPM-labeled CKIIpA. The experiments were performed as described in Figure 2 except that 10 µM CKIIp was included with CaM and Ca^2+^ in syringe A.

**Figure 4:**
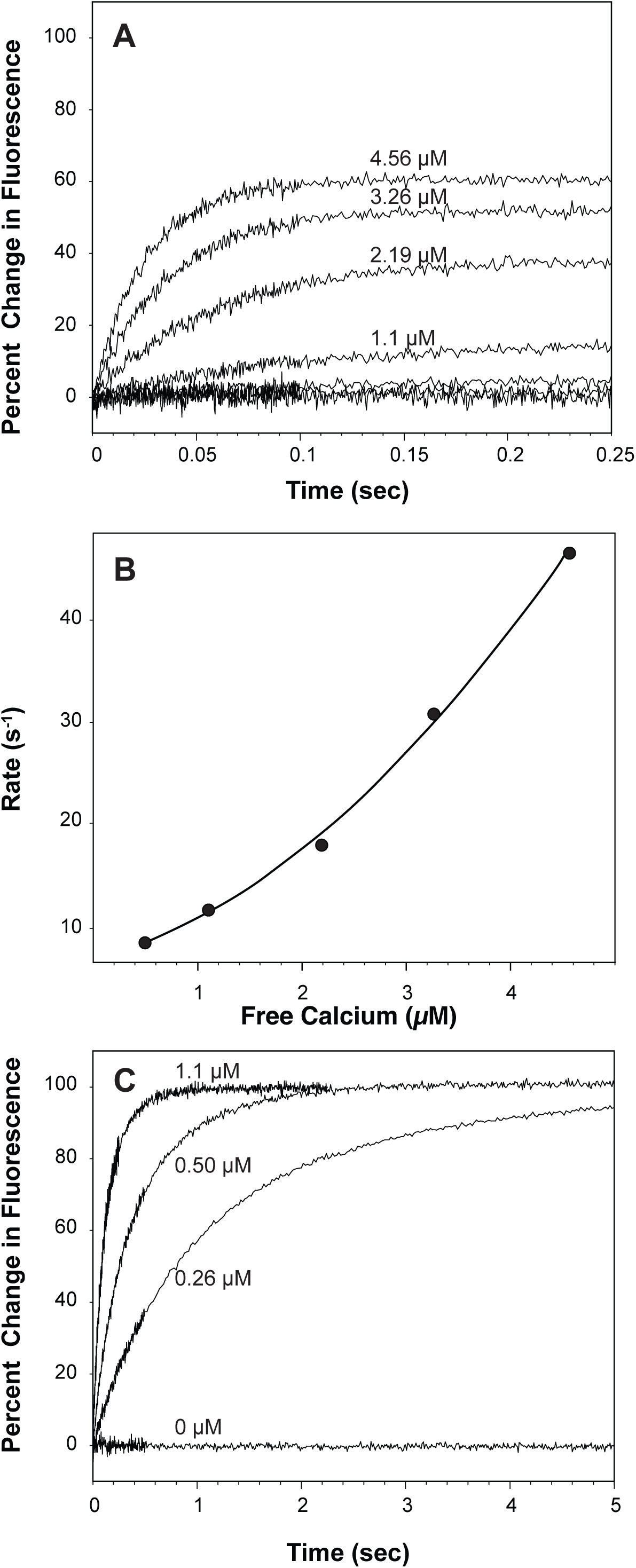
Rate of change of Tyr fluorescence from CaM upon association with Ca^2+^ in the absence or presence of CKIIp. *Panel A* shows the change in fluorescence when CaM (2.5 µM) was rapidly mixed with HEDTA solutions with Ca^2+^ buffered at 0, 0.26 µM, 0.50 µM 1.1 µM, 2.19 µM, 3.26 µM or 4.56 µM. No detectable change in Tyr fluorescence was observed at 0.26 µM Ca^2+^ . *Panel B* plots the rate of change of Tyr fluorescence (see Table 2) versus the Ca^2+^ . The line shows a fit to a quadratic equation. *Panel C* shows the same experiment as in *Panel A* except 5 µM CKIIp was added to the CaM solution.

We used a FRET assay to further confirm domain-specific assignment of Ca^2+^ dissociation rates in the CaM/CKIIp complex. The FRET donor was CaM(C110)D with an IAEDANS probe coupled to position 110 in the C-domain of CaM. The FRET acceptor was CKIIpA with DDPM coupled to a C-terminal Gly-Cys extension. Figure 3C and Table 1 show that mixing Ca^2+^-CaM(C110)D/CKIIpA with excess EGTA causes an initial decrease in fluorescence with a rate of 5.8 s-1, followed by a slower increase in fluorescence with a rate of 0.23 s-1. We conclude that the decrease in fluorescence is due to release of Ca^2+^ from the N-domain of CaM(C110)D causing a decrease in the average distance between the acceptor and donor probes. The slower increase in fluorescence is due to elimination of FRET as the C-domain of CaM(C110)D is released to fully dissociate the complex. Importantly, the rates observed using FRET are essentially identical to those collected using Quin-2, which indicates tight coupling between dissociation of Ca^2+^ and fast release of CKIIp from the N- or C-domain of CaM.

Table 1 shows that binding CKIIp to Ca^2+^-CaM decreases the rate of Ca^2+^ dissociation from the N- and C-domains about 250- and 35-fold, respectively. If CKIIp does not change the Ca^2+^ association rate, then the Kd for Ca^2+^ binding to the N- and C-domains in the CaM/CKII are both about 0.06 µM, which is consistent with the values reported by Peersen et al. (6).

### Measuring association rates at low levels of free Ca^2+^ (Ca^2+^) to mimic the intracellular milieu

Our general experimental design was to employ fluorescent reporters described above to monitor the overall rate of binding CKIIp to CaM after rapidly mixing apo CaM with solutions of CKIIp containing various levels of Ca^2+^ . A key feature of these experiments was to measure association rates at clamped Ca^2+^ levels of about 0.25 µM to 5 µM to mimic bulk cytoplasmic Ca^2+^ in response to stimuli (21). We postulated that this would allow us to detect formation of domain-specific encounter complexes, and to identify physiologically relevant effects of PEP-19 and Ng.

As described in detail in the Methods, sets of solutions were prepared using HEDTA (K_Ca_ ≈ 2.5 µM at pH 7.5) to buffer Ca^2+^. We used the Ca^2+^ chelators BAPTA and Br_2_BAPTA in control experiments to: 1) Verify the actual concentration of Ca^2+^ ; 2) Confirm that the equilibrium level of Ca^2+^ is not changed by binding Ca^2+^ to CaM; and 3) To confirm that replenishing the pool of Ca^2+^ is not a rate-limiting step, which is necessary to maintain constant Ca^2+^ during the time course of binding Ca^2+^ to CaM. Multiple experiments using multiple sets of Ca^2+^ buffers, ranging from 0.1 to 12 µM Ca^2+^, yielded consistent results. To allow ease of direct comparisons, the data presented in all Tables reported here were collected using a set of buffers with Ca^2+^ ranging from 0.26 µM to 4.56 µM.

Figure 4A and Table 2 show the rate of increase in Tyr fluorescence upon rapidly mixing free apo CaM with buffers of increasing concentrations of Ca^2+^ in the absence of CKIIp. The data best fit an exponential equation with a single rate at all levels of Ca^2+^ . Little change in fluorescence is observed at Ca^2+^ of 0.26 µM to 0.5 µM, and the change in fluorescence is about 84% of maximal at 4.6 µM Ca^2+^ . This is consistent with a K_Ca_ for the C-domain of free CaM of about 2 µM (6, 8, 22). Figure 4B shows a non-linear relationship between Ca^2+^ and the rate of change in Tyr fluorescence. We concluded previously that this was due to cooperative binding of Ca^2+^ to the C-domain of CaM (22). The data in Figure 4B was fit to a quadratic function to derive apparent pseudo-first order rates (k) at specific levels of Ca^2+^ using the first derivative. We selected Ca^2+^ of 1 µM (k) to compare pseudo-first order rates for data collected using fluorescence reporter assays since 1 µM is comparable to the elevated levels of Ca^2+^ in the bulk cytoplasm (21). Table 5 shows that the k_1.0_ derived from Tyr fluorescence experiments is 5.4 µM^-1^s^-1^. This is in good agreement with the association rate of 4.5 µM^-1^s^-1^ calculated for the C-domain using a K_Ca_ of 2 µM and k_off_ of about 9 s^-1^ from Table 1.

In striking contrast to free CaM, Figure 4C and Table 2 show that in the presence of CKIIp the increase in Try fluorescence approaches maximal even at the lowest Ca^2+^_F_. This is due to the large increase in Ca^2+^ binding affinity of CaM upon binding to CKIIp (6, 7). This means that the large change in fluorescence at low Ca^2+^_F_ reflects the rate of accumulation of the CKIIp/Ca^2+^-CaM complex as CKIIp binds to Ca^2+^-CaM and traps it in the Ca^2+^-bound form. The rate of change in Tyr fluorescence is significantly slower and more complex in the presence of CKIIp. The data in Figure 4C best fit an exponential equation with two rates up to 1.1 µM Ca^2+^, but only a single rate above 1.1 µM (see Table 2). The faster rate observed at all levels of Ca^2+^ is generally comparable in magnitude to the single rate observed for CaM in the absence of CKIIp. The implication of this with respect to the sequence of interactions between CKIIp and CaM are presented in the Discussion.

### Use of inter molecular FRET (inter-FRET) to monitor association rates

We used an inter-FRET assay with IAEDANS-labeled CaMD as a doner, and CKIIpA with DDPM coupled to a C-terminal GC extension as an acceptor. Unlike Tyr fluorescence, the inter-FRET assay is a more direct sensor of association between CaM and CKIIp and has the potential to sense binding events at either the N- or C-domain. Based on the crystal structure of CKIIp bound to Ca^2+^-CaM (19), we anticipated that the greatest inter-FRET effect would occur when the N-domain of CaM binds to the C-terminal portion of CKIIp, since this would allow the closest juxtaposition of donor and acceptor probes.

Figure 5A shows a robust inter-FRET response. Similar to Tyr fluorescence, the change in FRET approaches maximal amplitude even at 0.26 µM Ca^2+^_F_. Time-dependent changes in inter-FRET signals are complex, and best fit a tri-exponential equation with the slowest rate representing <7% of the overall change in amplitude. Interestingly, Supplemental Figure S1 shows that a slow, small and multicomponent change in the inter-FRET signal is seen even when CaMD was mixed with CKIIpA in the presence of 5 mM Ca^2+^ . These slow changes were not observed using Tyr fluorescence, and the amplitude of change observed at 5 mM Ca^2+^ was <10 percent of the total change in inter-FRET fluorescence observed at low Ca^2+^ . When mixed with 5 mM Ca^2+^, both the N- and C-domains of CaM would be saturated with Ca^2+^ within the dead-time of the fluorimeter, and binding of CKIIpA to Ca^2+^-CaMD would also occur in the dead time given the k_on_ of 120 µM-1s-1 (16) and concentrations of CKIIp and CaMD of 2.5 µM and 5 µM, respectively. Based on this, we conclude that the slow change in inter-FRET at 5 mM Ca^2+^ is due to a minor population of CKIIp that binds in a parallel orientation, and then undergoes an intramolecular reorientation, or must dissociate and rebind in the preferred antiparallel orientation (Step 5 in Figure 1). Since this minor slow rate is Ca^2+^ independent, our data analysis focused on rates corresponding to the two major components of change in inter-FRET fluorescence. As described in more detail in the Discussion our conclusion is that the dominant pathway for binding CaM to CKIIp involves formation of an encounter complex between the N-domain of CaM via step 2N in Figure 1.

**Figure 5:**
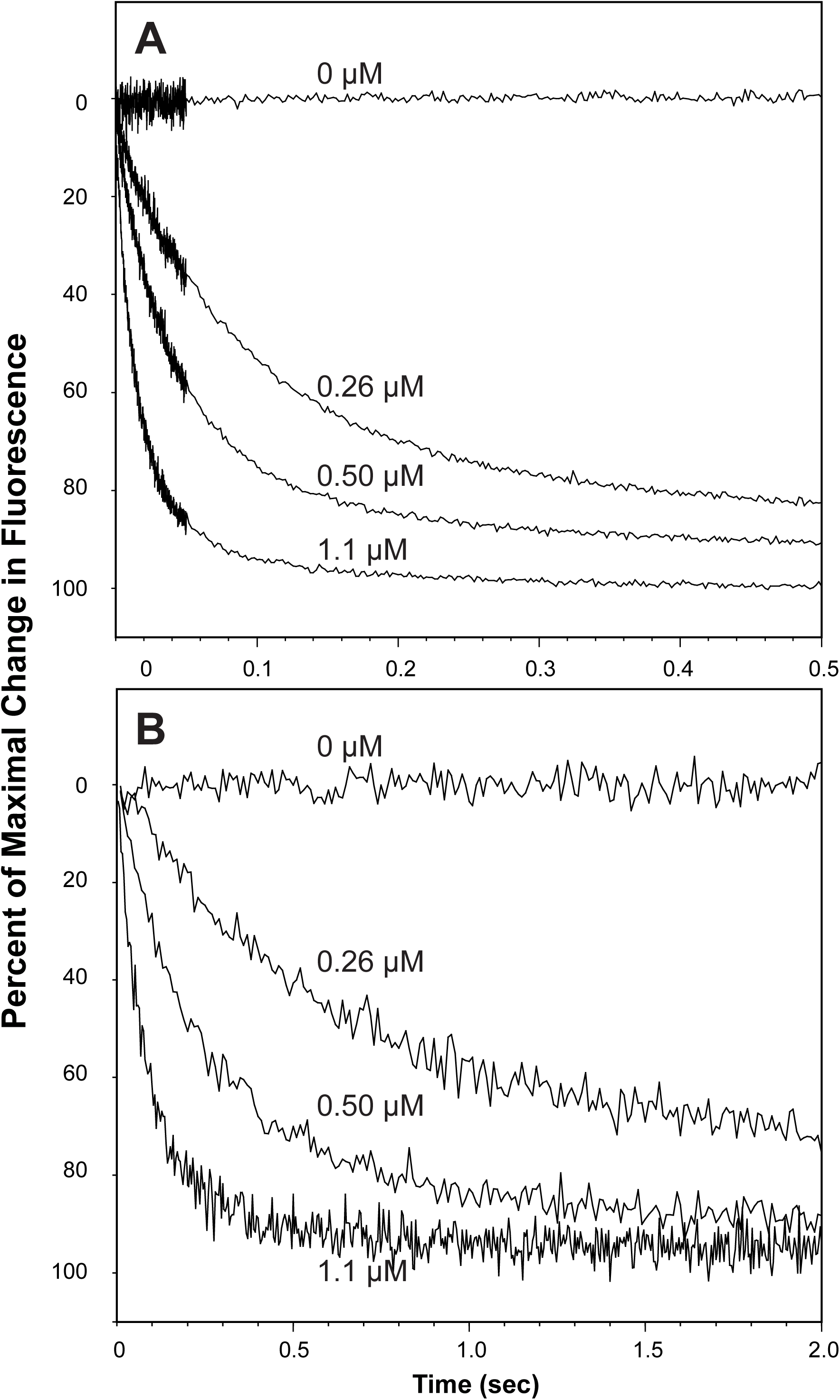
Use of inter and intra-FRET assays to monitor the rate of association of Ca^2+^-CaM with CKIIp: The inter-FRET assay in *Panel A* shows the change in fluorescence when a solution of CaMD (2.5 µM) and CKIIpA (5 µM) was rapidly was rapidly mixed with HEDTA solutions with the indicated concentrations of buffered Ca^2+^_F_. The intra-FRET assay in *Panel B* shows the change in fluorescence when a solution of CaMD/A (2.5 µM) and CKIIp (5 µM) was rapidly mixed with HEDTA solutions with increasing concentrations of buffered Ca^2+^ .

### Use of intra molecular FRET (intra-FRET) to monitor binding events

We also measured association rates using an intra-FRET assay that relies on FRET between donor and acceptor probes attached to the N- and C-domains of CaMD/A, which we used previously for assays performed under steady-state state conditions (17). Intuitively, intra-FRET would be most sensitive to Step 4 in Figure 1 since this involves a significant change in the relative distance between the N- and C-domains of CaM. Figure 5B shows the intra-FRET response at 0 to 1.1 µM Ca^2+^_F_. The data fit a single exponential equation, however, we cannot exclude the possibility that a second rate is obscured due to the low signal-to-noise ratio caused by a much lower probe-to-protein ratio of CaMD/A compared to CaMD. Table 2 shows that the single rate observed at all levels of Ca^2+^ are close in magnitude to the fast rate observed for Tyr fluorescence in the presence of CKIIp. The implications of these data with respect to the sequence of interactions between CKIIp and CaM are presented in the Discussion.

### Effects of PEP-19 on association of CKIIp with Ca^2+^-CaM

Figure 6 shows the typical effect of PEP-19 and Ng on response curves for Tyr, inter-FRET, and intra-FRET assays collected at 1.1 µM Ca^2+^, Tables 3 and 4 show rates derived at all levels of Ca^2+^ in the presence of PEP-19 or Ng, respectively. Figure 6A and Table 3 show that PEP-19 increased the rate observed using the Tyr fluorescence assay, and that two rates are observed at all levels of Ca^2+^ . The slow rate has the greatest amplitude at lower Ca^2+^, while the fast rate dominates at higher Ca^2+^ . The slow rate is similar in magnitude to the rate seen at all levels of Ca^2+^ in the absence of PEP-19. The pseudo first order rate constants in Table 5 show that in the presence of PEP-19, the fast rate (k_1.0R1_) of 85.4 µM^-1^s^-1^ is up to 16-fold greater than the two rates observed in the absence of PEP-19. Interestingly, Figure 6 and Tables 2 and 3 show that PEP-19 has little effect on the apparent rate of association of CKIIp with CaM when measured using either the inter-FRET or Inter-FRET assays. As described in more detail in the Discussion, these results point to a mechanism in which PEP-19 decreases cooperative Ca^2+^ binding to sites III and IV in the C-domain of CaM, which allows distinct stages of association between Ca^2+^-CaM and CKIIp with the C-domain. The slower stage is both required and rate-limiting for formation of the final compact complex between CaM and CKIIp.

**Figure 6:**
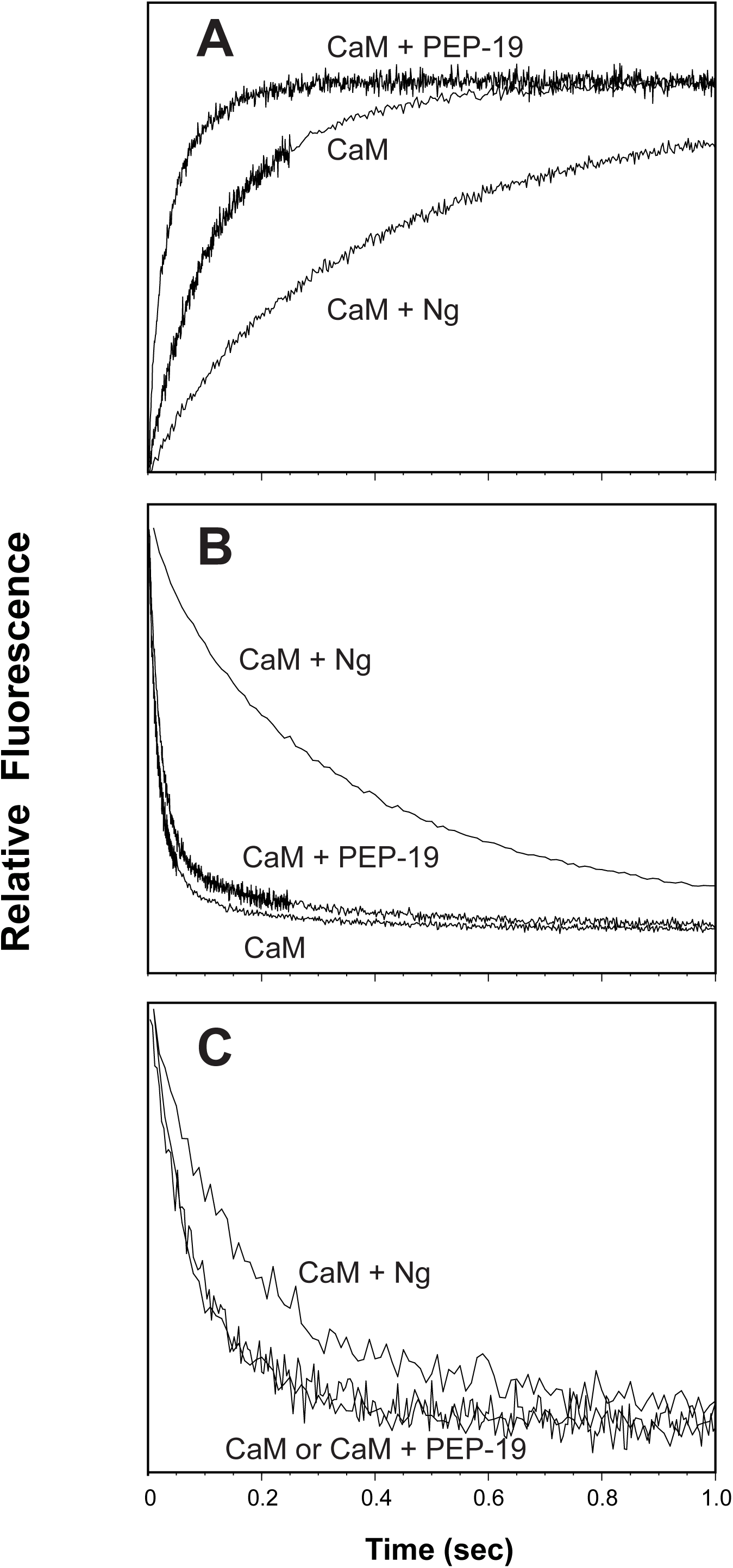
Effect of PEP-19 and Ng on the rate of association of CKIIp with Ca^2+^-CaM: The experiments were performed as described in Figures 4 and 5 except that the solutions of CaM and CKIIp in syringe A were prepared with or without 40 µM PEP-19 or 20 µM Ng. Syringe B contained an HEDTA solution with 1.1 µM. *Panels A, B and C* show the results using Tyr, inter-FRET and intra-FRET assays, respectively. The effects of PEP-19 and Ng on rate obtained at other concentrations of Ca^2+^ are reported in Tables 3 and 4.

**Table 3.** Rates of Association of CKIIp with Ca^2+^-CaM in the Presence of PEP-19.

| **Ca<sup>2+</sup>**<br>**(μM)** | **Tyrosine Fluorescence +PEP-19** |  |  |  | **Inter-FRET +PEP-19** |  |  |  | **Intra-FRET +PEP-19** |  |
| --- | --- | --- | --- | --- | --- | --- | --- | --- | --- | --- |
|  | Rate 1<br>(s<sup>-1</sup>) | Amp 1 | Rate 2<br>(s<sup>-1</sup>) | Amp 2 | Rate 1<br>(s<sup>-1</sup>) | Amp 1 | Rate 2<br>(s<sup>-1</sup>) | Amp 2 | Rate<br>(s<sup>-1</sup>) | Amp |
| 0.26 | 5.7 ± 0.5 | 33 | 1.6 ± 0.1 | 62 | 12 ± 0.8 | 25 | 3.8 ± 0.1 | 66 | 1.5 ± 0.1 | 79 |
| 0.50 | 17.9 ± 0.5 | 47 | 3.8 ± 0.6 | 55 | 19 ± 0.1 | 60 | 4.9 ± 0.8 | 35 | 3.9 ± 0.1 | 90 |
| 1.10 | 55 ± 4 | 47 | 14.4 ± 0.4 | 50 | 49 ± 0.6 | 86 | 7.2 ± 0.6 | 15 | 11.0 ± 0.3 | 100 |
| 2.19 | 158 ± 25 | 51 | 41 ± 2 | 56 | 136 ± 2 | 87 | 16.2 ± 1.2 | 15 | 30 ± 1 | 99 |
| 3.26 | 262 ± 26 | 73 | 58 ± 3 | 30 | 254 ± 7 | 81 | 32.7 ± 2.4 | 16 | 58 ± 4 | 80 |
| 4.56 | 346 ± 45 | 64 | 75 ± 5 | 20 | 414 ± 17 | 81 | 50.9 ± 3.6 | 18 | 99 ± 7 | 74 |

At least 5 stopped-flow traces were averaged and fitted using AppliedPhotophysics SX-Pro Data. The rates are shown with the estimated error of the fit.
To allow comparisons of amplitudes obtained from biphasic fits, as well as fits for different levels of Ca<sup>2+</sup><sub>F</sub>, the amplitudes are expressed relative to the overall amplitude at the Ca<sup>2+</sup><sub>F</sub> that gave the largest overall change in fluorescence within the data set.

**Table 4.**
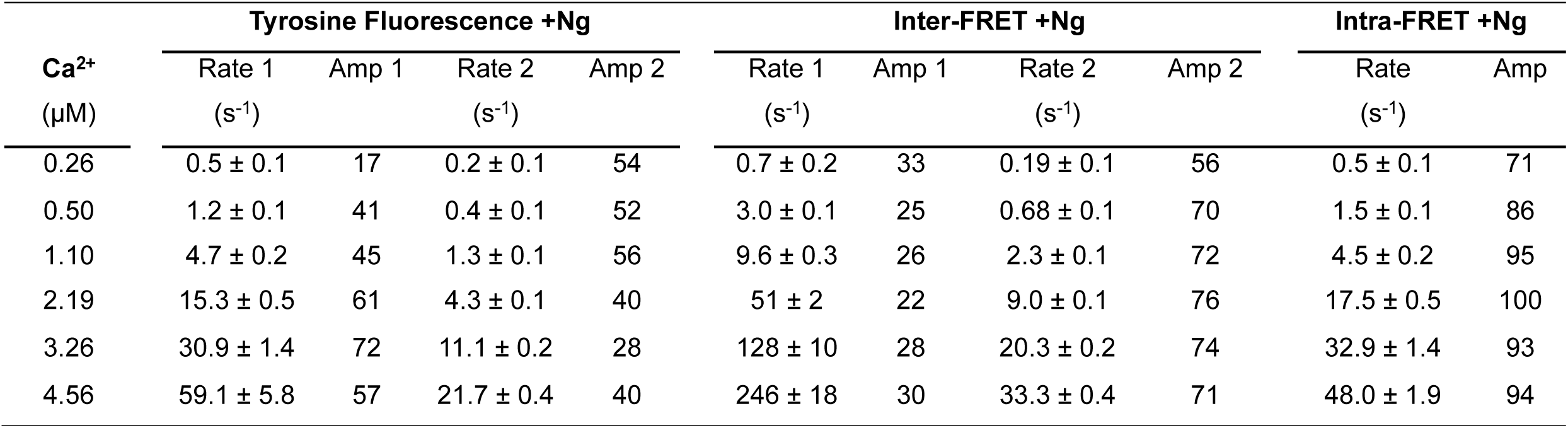
Rates of Association of CKIIp with Ca^2+^-CaM in the Presence of Ng.

### Ng decreases the rate of association of CKIIp with Ca^2+^-CaM

Figure 6 and Table 4 shows that Ng significantly decreases the overall rate of association of CKIIp with CaM when measured using all assays. This effect is most evident using the inter-FRET assay, and is most pronounced at lower levels of Ca^2+^ . It is reasonable to assume that the inter-FRET response with the fastest rate is due to formation of the preferred encounter complex along the pathway to final complex formation. Interestingly, the faster rate in the presence of Ng has a lower amplitude of fluorescence change relative to the absence of Ng, or the presence of PEP-19, which suggests a significant difference in the nature the encounter complex in the presence of Ng. The rates observed for the intra-FRET assay in the presence of Ng is very similar in magnitude at all levels of Ca^2+^ to the faster rate seen using Tyr fluorescence.

Data for the inter-FRET assay in Tables 2 and 4 show that the relative amplitudes of the fast rates dominate in the absence of Ng, but the amplitudes are greater for the slow rate in the presence of Ng. The magnitude of this effect is also seen from k_1.0_ values in Table 5. For example, Ng decreases the dominate k_1.0_ rate observed using the inter-FRET assay from 84.2 s^-1^µM^-1^ to 4.6 s^-1^µM^-1^.

Two rates are also detected by Tyr fluorescence in the presence of Ng at all levels of Ca^2+^ . Table 5 shows that the faster k_1.0_ of 6.4 s^-1^µM^-1^ measured using Tyr fluorescence is comparable to the slower inter-FRET rate of 4.6 s^-1^µM^-1^, and the single intra-FRET rate of 3.8 s^-1^µM^-1^. This suggests that these rates are due to the same kinetic step, and are likely due to the formation of a final compact complex between CaM and CKII.

**Table 5.** Pseudo First Order Calcium Dependent Association Rate Constants.

|  | Tyr | Tyr + CKIIp |  | Inter-FRET |  | Intra-FRET |
| --- | --- | --- | --- | --- | --- | --- |
|  | K<sub>1.0</sub> | K<sub>1.0R1</sub> | K<sub>1.0R2</sub> | K<sub>1.0R1</sub> | K<sub>1.0R2</sub> | K<sub>1.0</sub> |
| CaM | 5.4 | 10.6 | 5.2 | **84.2** | 8.0 | **15** |
| CaM + PEP-19 |  | 85.5 | 17.9 | **62.1** | 6.6 | **13.8** |
| CaM + Ng |  | 6.4 | 1.8 | 22.5 | 4.6 | **3.8** |

Rates (K) were derived by first fitting the data in Tables 1 to 3 to linear or quadratic functions. The best fits were used to calculate the first derivative (tangent) at 1.0 $\mu\text{M}$ $\text{Ca}^{2+}_{\text{F}}$ , which is the apparent pseudo first order rate constant at that level of $\text{Ca}^{2+}_{\text{F}}$ .
Numbers in bold indicate rates associated with the dominant amplitude.

### Ng and PEP-19 cause a nonlinear relationship between Ca^2+^ and the rate of initial complex formation

An interesting feature of Figure 4B is the nonlinear relationship between Ca^2+^ and the rate of Ca^2+^ binding to free CaM measured using Tyr fluorescence. We observed this previously (22) and attribute it to cooperative binding of Ca^2+^ to the C-domain of CaM. This led us to ask whether the rate of initial binding of Ca^2+^-CaM to CKIIp also exhibits nonlinearity with respect to levels of Ca^2+^ . Figure 7 shows the Ca^2+^ dependence of the fast rate observed using the inter-FRET assay, which is the best indicator of formation of the initial encounter complex. A linear relationship is seen for CaM. This is consistent with complex formation being primarily driven by the N-domain, which binds Ca^2+^ with less cooperativity than the C-domain. Interestingly, in the presence of PEP-19 and especially Ng, the curves are nonlinear, which indicates apparent positive cooperativity. The inset shows the ratio of the pseudo first order rate constants at 0.25 µM and 4.5 µM Ca^2+^ in the absence or presence of PEP-19 or Ng. A ratio of >20 in the presence of Ng means that the rate of initial complex formation is more responsive to Ca^2+^ at higher levels of Ca^2+^ .

**Figure 7:**
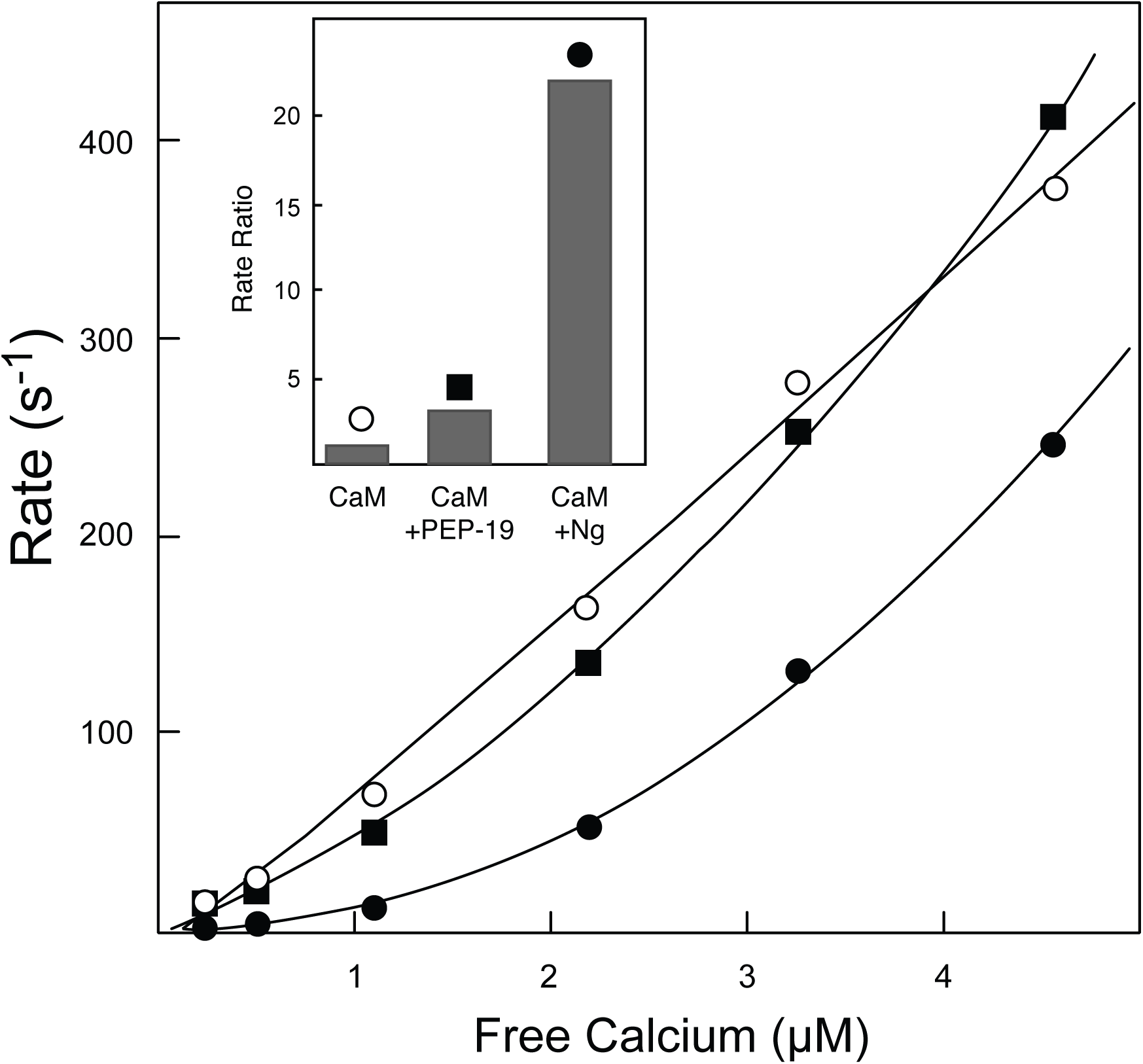
Ng and PEP-19 confer apparent cooperative binding of CKIIp to Ca2+-CaM: The curves show plots of Ca^2+^ versus the rates for the fast component (Rate 1) measured using the inter-FRET assay in the absence (○) or presence of PEP-19 (▪) or Ng (●). The inset shows the ratio of apparent pseudo first order rate constants at 4.5 µM and 0.25 µM Ca^2+^ (k /k).

### The Gly-rich C-terminal portion of Ng is required for its effect on association of CKIIp with CaM

The primary sequence of Ng has an IQ motif that is flanked on its N-terminal side by an acidic sequence, and on its C-terminal side by a Gly-rich sequence. Interestingly, we concluded using NMR that the Gly-rich C-terminal region of Ng associates with the N-domain Ca^2+^-CaM, while PEP-19, which lacks a Gly-rich region, binds only to the C-domain of CaM (14, 22). These observations led us to determine the effect of the Gly-rich region in Ng on the rate of association of CKIIp with Ca^2+^-CaM. Figure 8 shows that Ng(13–49), which includes both the acidic and IQ motifs, but is missing the Gly-rich C-term sequence, only slightly slows the rate of association of CKIIp with CaM. Thus, the Gly-rich region of Ng that interacts with the N-domain of CaM is primarily responsible for decreasing the rate of association of CKIIp with CaM.

**Figure 8:**
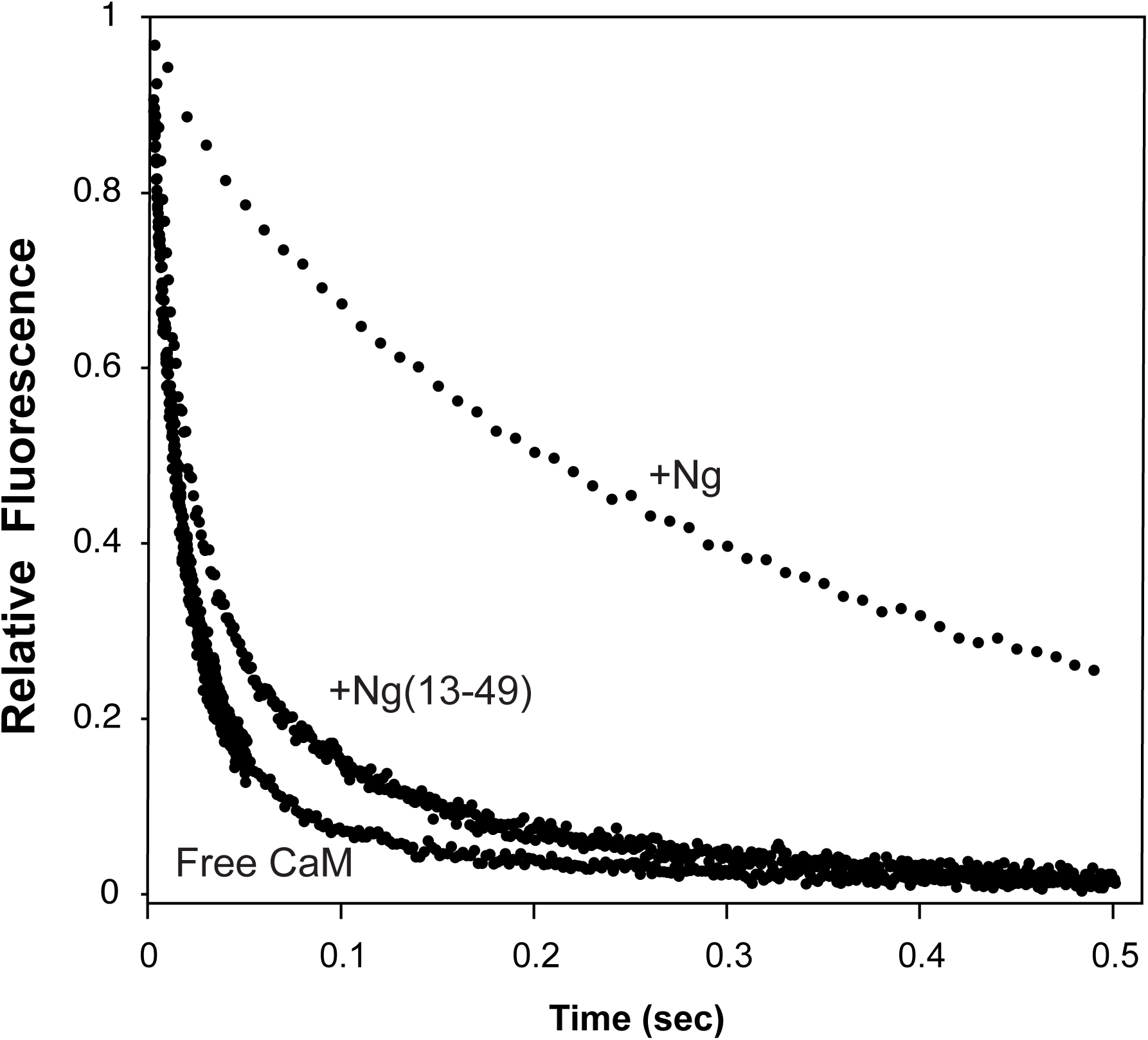
The effect of Ng on the rate of association of CKIIp with Ca^2+^-CaM is due to its C-terminal Gly-rich sequence: The inter-FRET experiments were performed as described in Figure 6B except that syringe A was prepared with or without 40 µM Ng or Ng(13–49). Syringe B contained an HEDTA solution with 1.1 µM Ca^2+^ .

### Ng slowly dissociates from Ca^2+^-CaM

We used NMR to determine if Ng or PEP-19 effectively competes with CKIIp for binding to Ca^2+^-CaM. Supplemental Figure 2 shows that binding CKIIp causes large chemical shift perturbations for almost all backbone amides in the ^1^H-^15^N-HSQC spectrum of ^15^N-lableled Ca^2+^-CaM. Supplemental Figures 3 and 4 show that addition of excess PEP-19 or Ng to the Ca^2+^-CaM/CKIIp complex caused little or no change to the HSQC spectra. This indicates that CKIIp, Ng and PEP-19 bind to the same or overlapping sites on Ca^2+^-CaM, and that association of CKIIp with Ca^2+^-CaM effectively prevents binding of Ng and PEP-19 under equilibrium conditions since its affinity for Ca^2+^-CaM is orders of magnitude greater than Ng or PEP-19.

Even though CKIIp will fully displace Ng or PEP-19 from Ca^2+^-CaM at equilibrium, the rate of dissociation of these modulators from Ca^2+^-CaM to expose binding sites for CKIIp could affect the rate at which equilibrium is achieved. This step does not seem to be rate limiting for PEP-19 since we used a FRET assay to show that PEP-19 fully disassociates from Ca^2+^-CaM within the dead time of the stopped-flow fluorimeter (23). Figure 9 shows a similar FRET experiment to monitor the dissociation of Ng from Ca^2+^-CaM. The data best fit an exponential equation with two rates of 90 s^-1^ and 15 s^-1^, with approximately equal amplitudes. Both rates are significantly slower than the rate of dissociation of PEP-19 from Ca^2+^-CaM, and are likely due to sequential dissociation Ng from the N- and C-domains of CaM. Together, these data show that Ng does not effectively compete with CKIIp for binding to Ca^2+^-CaM under steady state conditions, but could inhibit the rate of association due to its relatively slow rate of dissociation.

**Figure 9:**
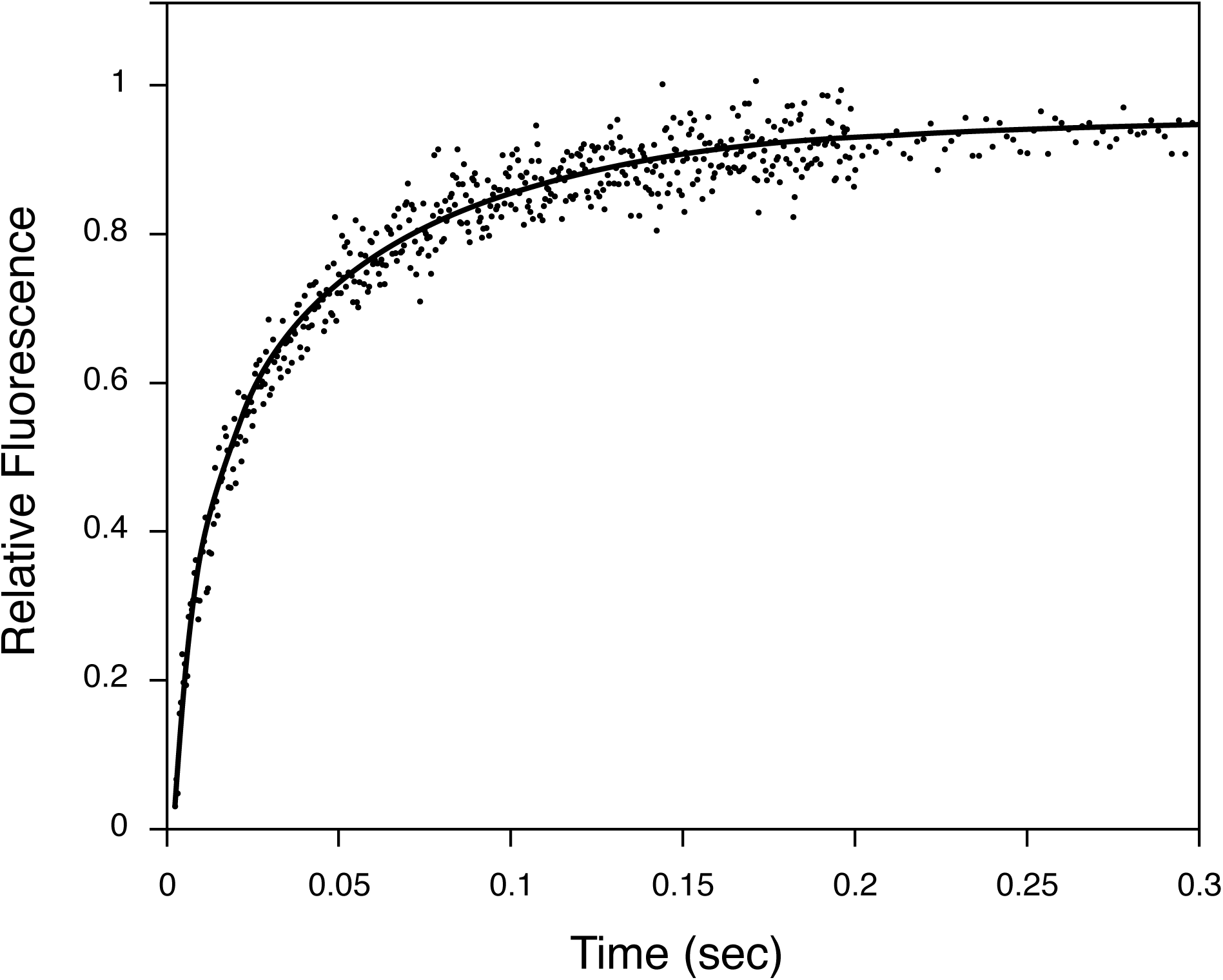
Rate of dissociation of Ng from Ca^2+^-CaM: Syringe A contained 1 μM IAEDANS-labeled CaM(D2C), and 10 μM NgD. Syringe B contained 20 μM unlabeled CaM. Both syringes contained 5 mM Ca^2+^. The increase in fluorescence reflects the rate of dissociation of labeled CaM from NgD and subsequent binding of excess unlabeled CaM.

## DISCUSSION

### Premise for the study

Binding Ca^2+^-CaM to target proteins can greatly increase its Ca^2+^ binding affinity due primarily to a decrease in the rate of dissociation of Ca^2+^ from the complex, and the magnitude of this effect is a function of the CaM binding sequence in the target (5, 6, 9, 10). This phenomenon likely contributes to the differential inactivation of CaM targets as Ca^2+^ levels fall. However, much less is known about the rate of Ca^2+^ dependent association of Ca^2+^-CaM with targets during a rise in Ca^2+^ levels, or during oscillations in Ca^2+^. In this study we used a target peptide derived from CaM Kinase II to provide a tractable model system to establish the mechanism and rate of association of both domains of Ca^2+^-CaM with a high-affinity target in the absence or presence of PEP-19 or Ng. Importantly, this was done at levels of free Ca^2+^ of around 1 µM to mimic the intracellular milieu, and to enhance the potential to observe functional consequences of divergent Ca^2+^-binding properties of the N- and C-domains of CaM.

### The N-domain of Ca^2+^ can drive association with CKIIp

Our first goal was to establish a baseline model for the sequence of Ca^2+^-dependent interactions between CaM and CKIIp in the absence of Ng or PEP-19, with a special focus on whether formation of the initial encounter complex was driven by the N- or C-domain of CaM. Early models focused on initial contact between the C-domain of CaM and target peptides along pathway 2C in Figure 1, primarily because the affinity of Ca^2+^ binding to the C-domain is greater than the N-domain (6, 7). However, we anticipated that initial contact involving the N-domain along pathway 2N could dominate since the rate of association of Ca^2+^ with the N-domain is much greater than the C-domain, and since Table 1 shows that the Ca^2+^ binding affinity of the N-domain is increased about 250-fold (K_Ca_ = 0.06 µM) upon binding to CKIIp due to a large decrease in the rate of Ca^2+^ dissociation. We reasoned that an encounter complex formed between CKIIp and the N-domain of CaM would be stabilized and accumulate due to this increase in the Ca^2+^ binding affinity. The following logic supports this hypothesis.

Domain-specific interactions between CaM and CKIIp can be deduced by comparing the rates of change of Tyr fluorescence, which is an indicator of Ca^2+^ binding to the C-domain, with the rates of change observed using the inter-FRET and intra-FRET assays, which are primary indicators of initial binding and final complex formation, respectively. If initial binding of CKIIp is driven primarily via the C-domain of CaM, then the dominant inter-FRET rate should be similar or slightly slower than the rate of Ca^2+^ binding to the C-domain, but this is not the case. The data in Table 2 show that the dominant rate (greatest amplitude) of complex formation observed using the inter-FRET assay is much faster at all levels of Ca^2+^ than the rate of association of Ca^2+^ with the C-domain of CaM determined using Tyr fluorescence in the presence of CKIIp. Moreover, the rate observed using the intra-FRET assay is comparable to the rate of binding Ca^2+^ to the C-domain in the presence of CKIIp. From this we conclude that the dominant pathway for complex formation is via Step 2N in Figure 1, and involves rapid binding of Ca^2+^ to the N-domain of CaM, followed by rapid binding of the N-domain to CKIIp and stabilization of the encounter complex due an increase in the Ca^2+^ binding affinity in the N-domain. This is followed by final complex formation along Steps 3C and 4C, which correlates with slower Ca^2+^ binding to the C-domain. This sequence of interactions could differ within the context of intact target proteins due to factors such as different accessibilities of binding sites for the N- and C-domains of CaM, nevertheless, it provides a well-characterized baseline kinetic model for interpreting the effects of PEP-19 and Ng.

### PEP-19 affects the cooperativity of Ca^2+^-binding to the C-domain of CaM

Table 2 shows that association of Ca^2+^-CaM with CKIIp results in a biphasic change of inter-FRET fluorescence. Table 3 shows that the inter-FRET rates and amplitudes are generally similar in the absence or presence of PEP-19 with the faster rate having the dominant amplitude, especially at higher levels of Ca^2+^_F_. Thus, PEP-19 does not greatly affect the dominant pathway for complex formation via Step 2N in Figure 1. This is not surprising since PEP-19 binds selectively to the C-domain (13), and we concluded above that initial contact with CKIIp is primarily driven by the N-domain. We were also not surprised that PEP-19 increased the Ca^2+^-dependent rate of Tyr fluorescence in the presence of CKIIp (see Figure 6A) since PEP-19 increases the rate of Ca^2+^ binding to the C-domain (13). However, we anticipated that PEP-19 would increase the rates observed using the intra-FRET or inter-FRET assays, but this was not seen. Modulation of cooperative Ca^2+^ binding to the C-domain may provide a mechanism to explain these results. We showed previously that PEP-19 decreases the cooperativity of Ca^2+^ binding to the C-domain (22). The data in Table 3 are consistent with this observation since Tyr fluorescence detects two events with fast and slow rates at all levels of Ca^2+^_F_ in the presence of PEP-19 and CKIIp. Interestingly, the slow rate of change in Tyr fluorescence in the presence of PEP-19 (Rate 2 in Table 3) is similar in magnitude to both the fast rate in the absence of PEP-19 (Rate 1 in Table 2), and with the rates observed using inter-FRET in the absence or presence of PEP-19. These data support a model in which PEP-19 promotes rapid binding of Ca^2+^ to site III or IV in the C-domain, but that the 1-Ca^2+^ form of the C-domain is not competent to bind to CKIIp, or binds in a conformation that does not alter Tyr fluorescence. Slower Ca^2+^ binding to the second site is required for the C-domain to associate with CKIIp, and is detected by a second phase of change in Tyr fluorescence, as well as a change in the inter-FRET signal. Based on the NMR solution structure of PEP-19 bound to apo-CaM, we propose that Ca^2+^ binds first to Site III and then Site IV (24). The rapid rate of dissociation of PEP-19 from the C-domain of Ca^2+^-CaM (>400 s-1) (23) is consistent with this model since PEP-19 does not present a kinetic barrier that would inhibit or slow the rate of binding of CKIIp.

### Ng greatly decreases the rate of association of CKIIp with Ca^2+^CaM

In contrast to PEP-19, all assays show that Ng significantly decreases the overall rate of association of CKIIp with CaM, and that this effect is especially striking at lower concentrations of Ca^2+^ . For example, Table 5 shows that Ng decreases the dominant k_1.0_ rate measured using inter-FRET by almost 20-fold from 84.2 s^-1^µM^-1^ to 4.6 s^-1^µM^-1^. We conclude from Figures 8 and 9 that this property of Ng is due to the combined effects of two factors. First, the Gly-rich C-terminal portion of Ng binds to the N-domain of CaM to inhibit interaction between the N-domain and CKIIp. Second, Ng dissociates relatively slowly from Ca^2+^-CaM to further inhibit the rate of binding to CKIIp. These two factors suggests that Ng directs CKIIp to bind initially to the C-domain by sterically blocking binding the CKIIp binding site in N-domain. This would explain why the dominant amplitude seen for the inter-FRET assay in the presence of Ng is associated with the slower rate (Amp 2 in Table 4), instead of the faster rate (Amp 1 in Table 2) in the absence of Ng. It is also possible that the preferred pathway in the presence of Ng depends on the level of Ca^2+^ . For example, pathway 2C may be preferred at lower Ca^2+^, while 2N dominates at higher Ca^2+^ . This would be consistent with the fact that Ng decreases the Ca^2+^ binding affinity of CaM (14). Regardless of the precise mechanism, it is clear that Ng greatly slows the rate of binding CKIIp to Ca^2+^-CaM, and could differentially modulate the rate of activation of CaM target proteins in cells.

### Ng tunes the Ca^2+^ sensitivity of complex formation

Figure 7 shows that PEP-19, and especially Ng, causes a nonlinear relationship between Ca^2+^_F_ and the rate of association between CaM and CKIIp. Essentially, the Ca^2+^-sensitivity of CaM binding to CKIIp is decreased at low levels of Ca^2+^_F_, and there is a disproportionate increase in the rate of association at higher levels of Ca^2+^_F_. This may be due to multiple factors, including different pathways to complex formation at low versus high Ca^2+^_F_. Regardless of the precise mechanism, the data demonstrate that Ng has the potential to modulate CaM signaling by tuning the apparent Ca^2+^-sensitivity of complex formation. The extent of this phenomenon on the spectrum of CaM target proteins will likely depend on the mode of CaM binding, but Ng could, for example, provide a kinetic proof-reading mechanism to prevent unwanted activation of CaM-targets under oscillating but sub-threshold Ca^2+^-increases.

### Potential effects on distribution of CaM among targets

Intracellular levels of CaM are limiting relative to the total number of CaM binding proteins (25). This condition establishes a competitive scenario in which the largest fraction of Ca^2+^-CaM will associate with the highest affinity targets at equilibrium. However, equilibrium with respect to Ca^2+^ is an elusive intracellular condition due to frequent oscillations in Ca^2+^ levels, and computational models predict that oscillating Ca^2+^ levels are sensed by the Ca^2+^ binding properties of CaM to achieve differential activation of target proteins (26). Shuffling of CaM between targets during Ca^2+^ oscillations will depend on the rate of association of Ca^2+^-CaM with target proteins, and the rate of dissociation of Ca^2+^ from Ca^2+^-CaM/target complexes. The rate of association of Ca^2+^-CaM with targets will depend on factors such as local levels of and rate of diffusion of Ca^2+^, the local levels of free or bound CaM, and the accessibility of CaM binding sites on targets. The data presented here demonstrate that Ng can play a role in this process by significantly decreasing the rate of association of targets with Ca^2+^-CaM. This adds a new mechanism to modulate the distribution of CaM between targets during Ca^2+^ oscillations, one that can be regulated by phosphorylation of Ng, which inhibits binding to CaM (27).

## CONCLUSION

Ng and PEP-19 are small, intrinsically disordered proteins with no known biological activity other than binding to CaM, yet they have the potential to regulate CaM signaling pathways via multiple mechanisms. One mechanism is their ability to modulate the Ca^2+^ binding properties of CaM (13, 15). They bind CaM with relatively low affinity, and would be poor steady-state competitive inhibitors of high-affinity intracellular CaM targets, but CaM has multiples modes of interaction with targets, so Ng and PEP-19 could function as inhibitors of low-affinity targets that bind primarily to the N- or C-domain of CaM. They could also modulate the activation/inactivation of targets such as the voltage gated Ca^2+^ channels on which the N- and C-domains of CaM shift among different binding sites (28). The data presented here demonstrate that inhibiting the rate of association of Ca^2+^-CaM with high-affinity targets is another mechanism by which Ng can impact CaM signaling. The magnitude of this effect will be will likely depend on the mode of interaction of Ca^2+^-CaM with different targets, which could contribute to tuning the differential activation of targets in a complex cellular milieu where multiple targets compete for a limited supply of CaM. This mechanism could also influence the redistribution of CaM, and the coordinated activation of CaM targets during a train of Ca^2+^ oscillations.

## ABBREVIATIONS

CaM: calmodulin
CKIIp: amino acids 292-312 from calmodulin dependent protein kinase II
Ng: neurogranin
CaMD: CaM(K75C) labeled with IAEDANS

## APPENDICES

### AUTHOR CONTRIBUTIONS

J. Putkey designed the experiments, helped collect stop-flow data, prepared the figures and tables and wrote the manuscript. L. Hoffman and V. Berka help collect stop-flow data. X. Wang collected and processed the NMR data. All authors reviewed the manuscript.

### DECLARATON OF INTERESTS

The authors declare no competing interests.

## ACKNOWLEDGEMENTS

The authors wish to thank Dr. M. Neal Waxham for reviewing this manuscript and providing insightful comments. This work was supported in part by grant R01 GM104290 from the National Institutes of Health to J.A.P.

---

**Figure S1.**
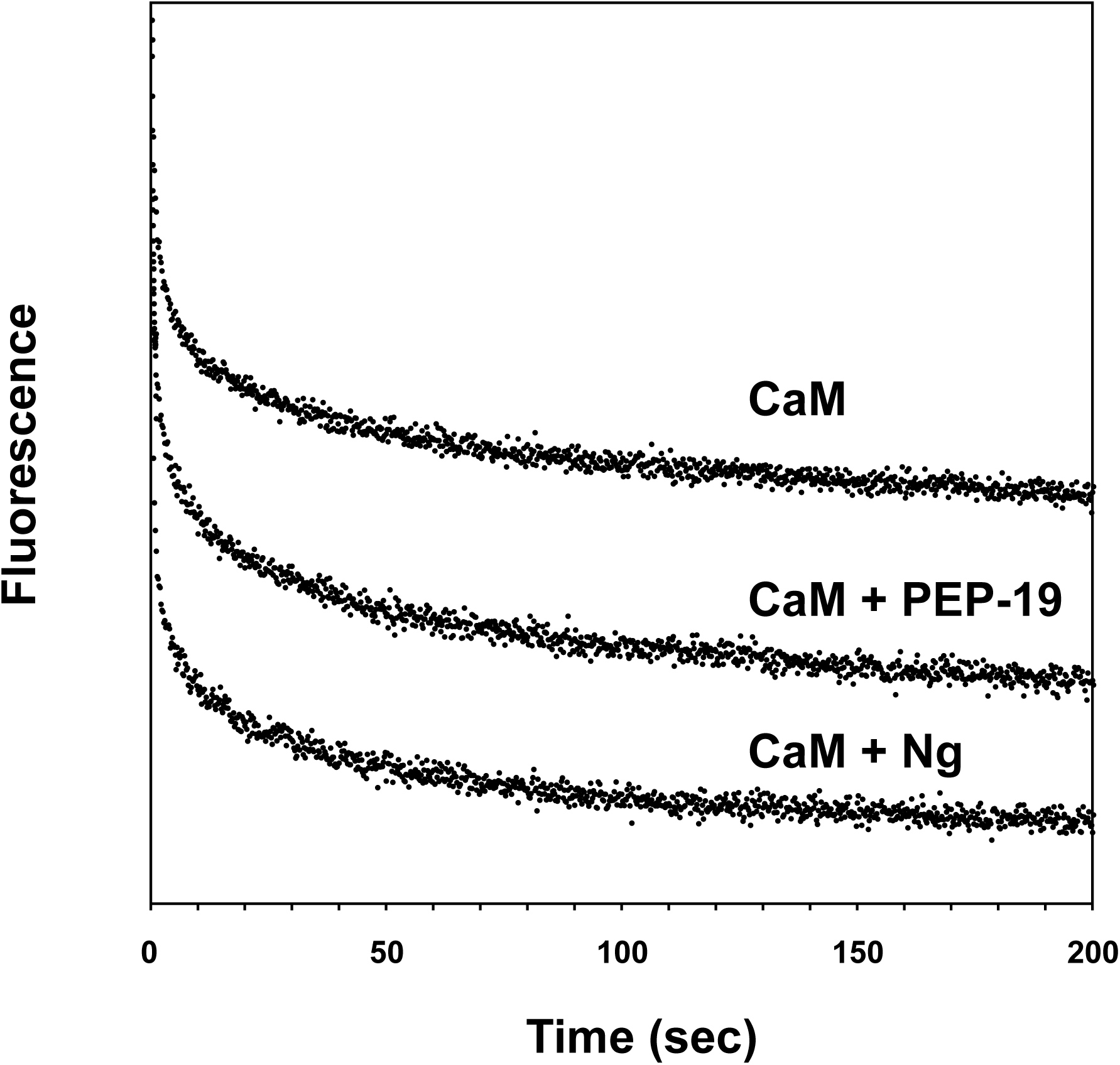

**Figure S2.**
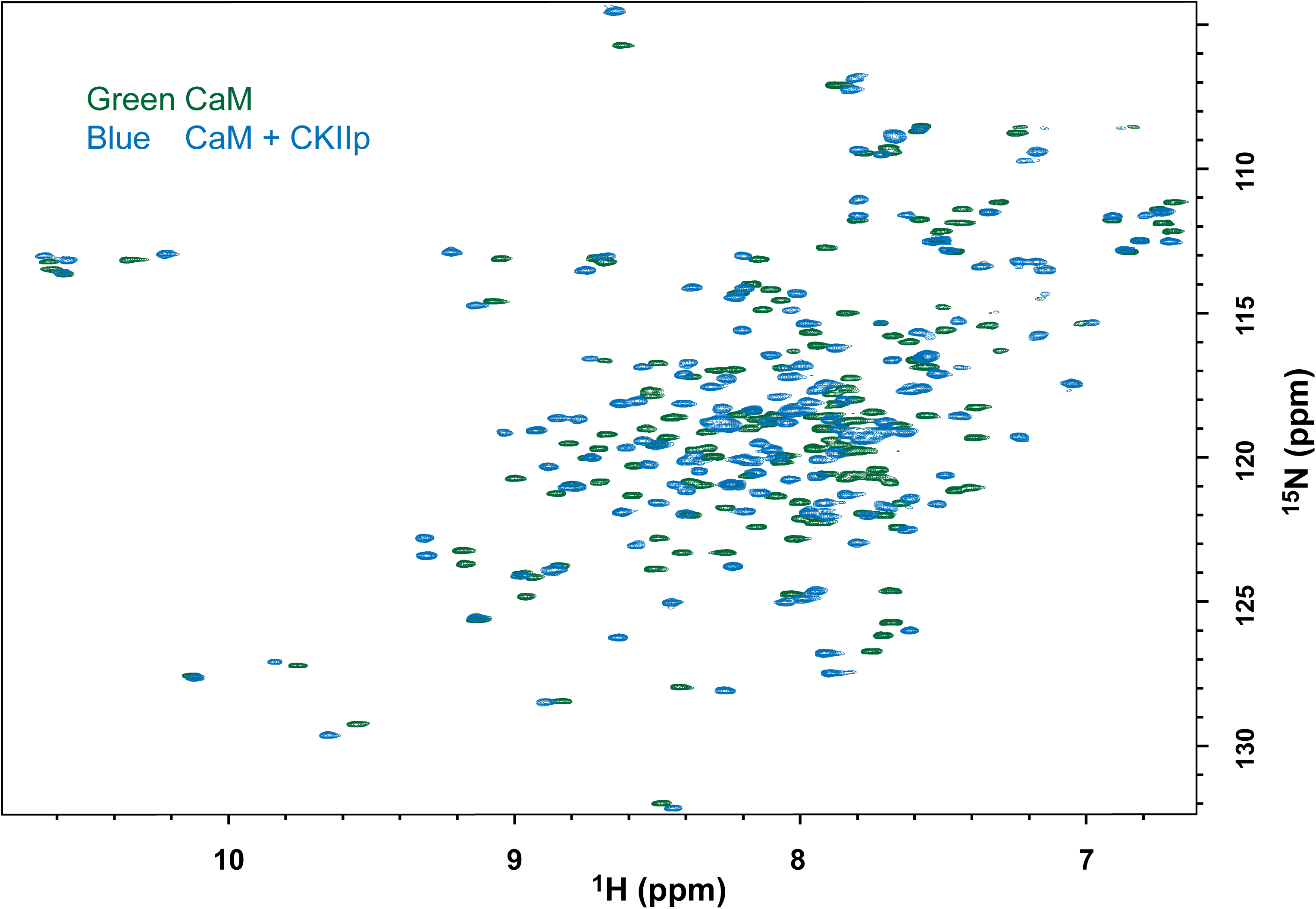

**Figure S3.**
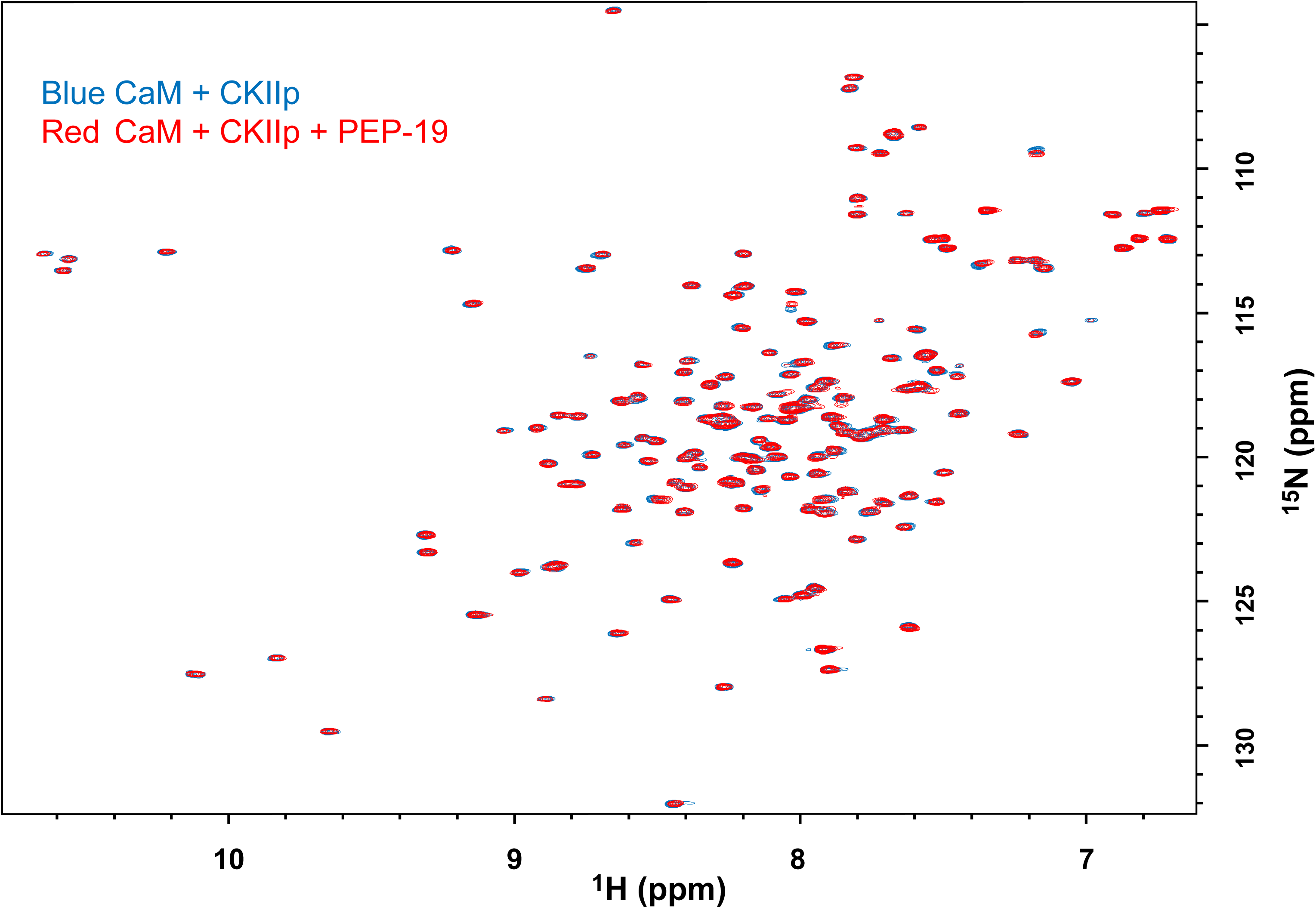

**Figure S4.**
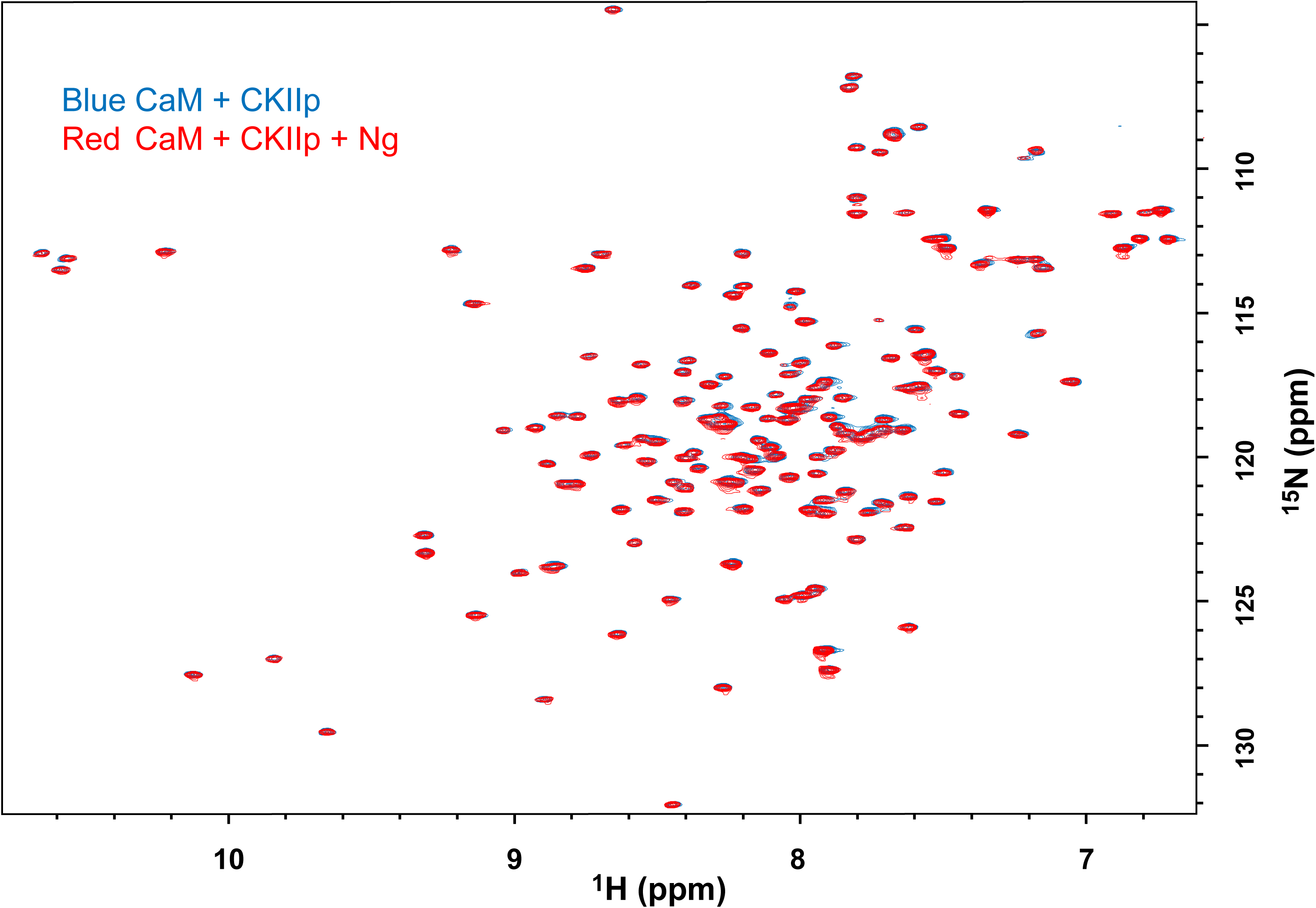

